# STRAIGHT-IN Dual: a platform for dual, single-copy integrations of DNA payloads and gene circuits into human induced pluripotent stem cell

**DOI:** 10.1101/2024.10.17.616637

**Authors:** Albert Blanch-Asensio, Deon S. Ploessl, Sara Cascione, Benjamin B. Johnson, Myrthe R. M. Berndsen, Nathan B. Wang, Valeria Orlova, Anna Alemany, Christine L. Mummery, Kate E. Galloway, Richard P. Davis

## Abstract

Targeting DNA payloads into human (h)iPSCs involves multiple time-consuming, inefficient steps that must be repeated for each construct. Here, we present STRAIGHT-IN Dual, which enables simultaneous, allele-specific, single-copy integration of two DNA payloads with 100% efficiency within one week. Notably, STRAIGHT-IN Dual leverages the STRAIGHT-IN platform to allow near-scarless cargo integration, facilitating the recycling of components for subsequent cellular modifications. Using STRAIGHT-IN Dual, we investigated how promoter choice and gene syntax influence transgene silencing, demonstrating the impact these design features have on reporter gene expression and forward programming of hiPSCs into neurons, motor neurons, and endothelial cells. Furthermore, we designed a grazoprevir-inducible synZiFTR system to complement the widely used tetracycline-inducible system, providing independent, tunable, and temporally controlled expression of different transcription factors within the same cell. The unprecedented efficiency and speed with which STRAIGHT-IN Dual generates homogenous genetically engineered hiPSC populations represents a major advancement for synthetic biology in stem cell applications and opens opportunities for precision cell engineering.

## Introduction

The efficient and precise insertion of DNA payloads into mammalian genome remains a major challenge for synthetic biology, disease modeling, and cell-based therapies. Applications ranging from biosensor and reporter integration to modelling disease variants, generating humanized mouse models, and programming cell fate all require reliable genomic integration tools (Balmas et al., 2023; Blanch-Asensio et al., 2022; Brandão et al., 2020; Hosur et al., 2022; Low et al., 2022; Pawlowski et al., 2017; M. Zhang et al., 2021). Recently, hybrid platforms combining genome editing technologies such as prime editing with site-specific recombinases (SSRs) have enabled the insertion of DNA fragments up to 36 kb, offering a promising route for complex genome engineering (Anzalone et al., 2022; Pandey et al., 2025; Yarnall et al., 2023). However, CRISPR/Cas9-based methods often suffer from off-target effects and variable efficiencies, necessitating extensive clonal screening.

SSR-based methods provide an alternative approach, enabling efficient and high-fidelity integration of large DNA payloads (>100 kb) into engineered landing pad (LP) cassettes (Blanch-Asensio et al., 2022; Brosh et al., 2021; Mitchell et al., 2021; Pinglay et al., 2022). Among these, the serine recombinase Bxb1 is particularly effective due to its high and irreversible integration efficiency, tolerance of large cargoes, and absence of pseudo-recognition sites in the human genome (Duportet et al., 2014; Hosur et al., 2022; Roelle at al., 2023). However, current SSR-based platforms are typically limited to single-copy integration at a single locus. While some platforms permit integration of two DNA payloads (Rosenstein et al., 2024), few support precise, allele-specific insertion of multiple transgenes. Alternative methods, such as cassette exchange which relies on two recombination sites (Matreyek et al., 2017), remain relatively inefficient.

Previously, we developed STRAIGHT-IN, a modular Bxb1-based system for single-copy transgene integration into human induced pluripotent stem cells (hiPSCs) (Blanch-Asensio et al., 2022). Although we also evaluated the recombinase ϕC31 for orthogonal integration of two donor plasmids, its low efficiency in hiPSCs precluded practical use. Recent mechanistic studies of Bxb1 recombination have engineered orthogonal *attP*/*attB* sites by altering the central “GT” dinucleotide to “GA”, which maintained high integration efficiency with minimal crossover between the recombination sequences (Jusiak et al., 2019; Roelle et al., 2023).

Building on these developments and aiming to address limitations of previous systems, we developed STRAIGHT-IN Dual, a hiPSC acceptor line containing two orthogonal Bxb1-compatible LPs. Using a single recombinase, this system enables simultaneous, and allele-specific integration of two DNA payloads into distinct alleles of the *CLYBL* locus, a genomically permissive site that supports stable, long-term transgene expression without perturbing endogenous genes (Cerbini et al., 2015). STRAIGHT-IN Dual also supports seamless removal of auxiliary sequences (e.g. selection markers and vector backbones), using the tyrosine recombinases Cre and Flp. This results in markerless and near-scarless integration of the DNA payload (Blanch-Asensio et al., 2022; Brosh et al., 2021; J. Li et al., 2020), reducing transgene silencing and facilitating iterative genomic modifications.

This dual LP platform facilitates scalable synthetic and stem cell biology studies while controlling for locus-specific and neighboring transgene effects. By enabling single-copy, allele-specific integration of distinct payloads, STRAIGHT-IN Dual supports systematic investigation of how promoter choice, gene syntax (i.e. the relative order and orientation of genes), and other regulatory elements influence transgene expression and stability across pluripotent and differentiated cell states. Moreover, we demonstrated that multiplexed integration of a panel of constructs allows single-pot characterization and identification via Sort-Seq (O’Connell et al., 2023), establishing a scalable framework for high-throughput screening of genetic components, including promoter libraries and synthetic gene circuits.

We further used STRAIGHT-IN Dual to identify optimal strategies for forward programming of hiPSCs into induced neurons (iNs), endothelial cells (iECs), and motor neurons (iMNs), demonstrating that distributing transcription factors across the two LPs enhanced differentiation outcomes. To expand transgene control capabilities, we adapted a grazoprevir-inducible synthetic zinc finger transcription regulator (synZiFTR) system (H.-S. Li et al., 2022) for use in hiPSCs. Combined with the doxycycline-inducible all-in-one Tet-On 3G system, this allowed precise and independent control of dual genetic programs. Notably, we used this to establish dual-fate cell lines in which we could specify distinct fates from a single cell line by applying different inducers, thereby opening opportunities for spatial patterning of multiple cell types from a genetically uniform background.

Taken together, STRAIGHT-IN Dual offers a robust and flexible platform for precise, multi-transgene integration and inducible control in hiPSCs. These capabilities make it uniquely suited for sophisticated cell engineering applications ranging from high-throughput genetic screening to combinatorial circuit design and multi-lineage modeling.

## Results

### Bxb1 mediates allele-specific integration of two DNA payloads in *CLYBL*

Building upon our previously developed STRAIGHT-IN acceptor hiPSC line that contained a single LP in one allele of *CLYBL* (Blanch-Asensio et al., 2023), we targeted the unedited *CLYBL* allele to establish a dual STRAIGHT-IN acceptor hiPSC line. The original STRAIGHT-IN LP line (LU99*_CLYBL*-bxb-v2) contains a bxb1-GT *attP* site for allele-specific targeting with GT donor plasmids, as well as an excisable fluorescent reporter (*EBFP2*) driven by the PGK promoter and a selection cassette (*BleoR*) designed as a promoter trap due to the lack of an ATG initiation codon and promoter sequence. The 5’ and 3’ ends of the LP are flanked by two heterospecific *loxP* and *lox257* sites, enabling Cre recombinase (Cre) to excise the auxiliary sequences both 5’ and 3’ of the payload following integration (**Figure 1a**).

**Figure 1.**
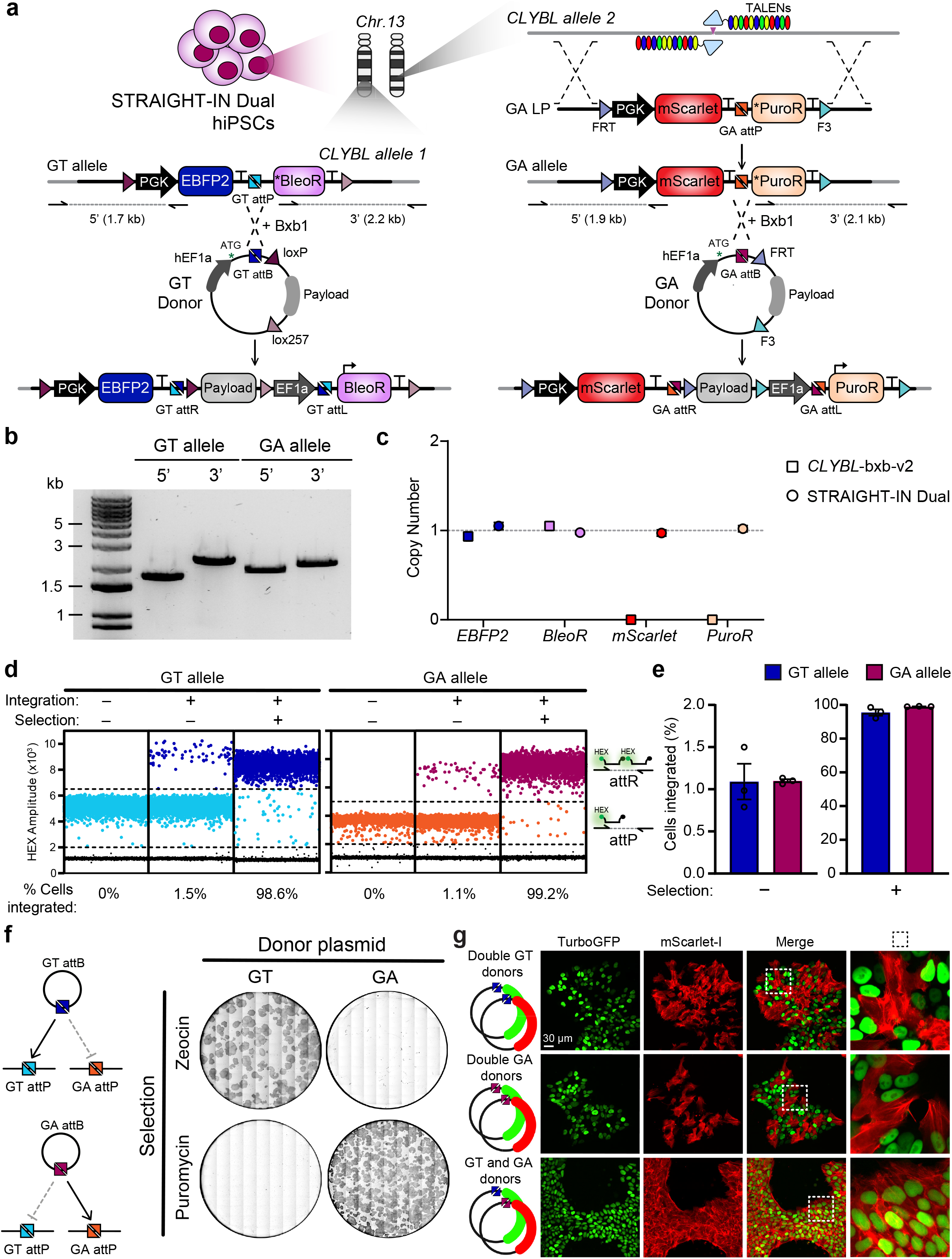
Targeted allele-specific integration of bxb1-GT and bxb1-GA payloads in the STRAIGHT-IN Dual hiPSC line. (***a***) Schematic of TALEN-mediated targeting of the GA-LP cassette into the second allele of the citrate lyase beta- like (*CLYBL*) locus and Bxb1 recombinase-mediated integration of GT and GA donor plasmids into their cognate LPs. Expression of the antibiotic resistance markers is activated only upon correct donor plasmid integration, which supplies the missing initiation codon (*). Half arrows indicate junction PCR primer sites. “Payload” indicates the location of the desired DNA cargo for targeting in both the donor plasmid and following genomic integration. “T” denotes polyadenylation sequences. (***b***) Junction PCR analysis confirming targeting of GT and GA LPs in the *CLYBL* locus. (***c***) ddPCR validating single-copy genomic integration of each LP cassette. The *CLYBL*-bxb-v2 hiPSC line serves as a control. Error bars, Poisson 95% CI. (***d***) Representative ddPCR dot plots showing GT and GA donor plasmid integration before and after antibiotic selection. Dots represent droplets containing the indicated sequence (*attR* or *attP*), with percentages showing the calculated integration efficiencies. (***e***) Mean integration efficiencies of GT and GA donor plasmids before (-) and after (+) antibiotic selection. N=3 independent transfections; error bars, ±SEM. (***f***) Schematic illustrating Bxb1-GT/GA recombination specificity (*left*), and alkaline phosphatase staining (*right*) confirming orthogonal payload integration at the GT and GA LPs.

Using TALENs, we targeted a novel LP containing a bxb1-GA *attP* site into the unedited *CLYBL* allele, thereby permitting allele-specific targeting with GA donor plasmids (**Figure 1a**). This bxb1-GA LP also includes a distinct fluorescent reporter (*mScarlet*), selection cassette (*PuroR*) and heterospecific *FRT* and *F3* sites recognized by Flp recombinase. We characterized one of the resulting hiPSC clones (STRAIGHT-IN Dual) expressing mScarlet by genotyping PCR, ddPCR and Sanger sequencing, confirming a single copy of the novel LP in the previously unedited *CLYBL* allele with no mutations in the recombination sites (**Figure 1b,c** and **Supplementary Figure 1a**). Additionally, the genomic integrity and normal karyotypic status of the hiPSC line was confirmed by ddPCR and G-band karyotyping (**Supplementary Figure 1b,c**). The hiPSC clone also expressed pluripotency markers and could differentiate into all three germ layers (**Supplementary Figure 1d-f**).

Next, we designed donor plasmids containing either *attB*-GT or *attB*-GA recombination sites. In the presence of Bxb1, these donor plasmids specifically integrate into their cognate LPs – *attB*-GT into the *attP*-GT allele and *attB*-GA into the *attP*-GA allele – allowing precise allele-specific cargo integrations (**Figure 1a**). Each donor plasmid contains the hEF1a promoter sequence and an ATG initiation codon that completes the promoter trap upon integration, leading to the expression of the respective selection markers (BleoR for the GT allele, PuroR for the GA allele) and conferring resistance to zeocin or puromycin.

We first evaluated the efficiency of GT and GA donor plasmid integration, observing comparable levels for both alleles before selection (1.09% ± 0.21 for GT, 1.10% ± 0.02 for GA; **Figure 1d,e**). Subsequent antibiotic selection led to robust enrichment of hiPSCs containing either the GT or GA donor plasmids (95.52% ± 1.87 for GT, 99.06% ± 0.08 for GA; **Figure 1d,e**). Additionally, co-transfection of both donor plasmids and a Bxb1 expression plasmid, followed by dual antibiotic selection, resulted in >93.9% of hiPSCs containing both integrated payloads (**Supplementary Figure 11,g**).

To confirm allele specificity, we independently transfected either the GT or GA donor plasmids and subjected cells to selection with the non-matching antibiotic. If the GT donor incorrectly integrated into the *attP*-GA LP, puromycin resistance would be conferred. Likewise, incorrect GA donor integration into the GT LP would lead to zeocin resistance (**Figure 1f**). In both cases, selection resulted in complete cell death, confirming the orthogonality of the alleles (**Figure 1f**).

To further validate the orthogonality of the GT and GA alleles, we cloned green and red fluorescent reporters (*TurboGFP* and *mScarlet-I*, respectively) into both donor plasmids (**Figure 1g**). *TurboGFP* was tagged with a nuclear localization sequence (NLS), and *mScarlet*-I with an actin-targeting signal, enabling unambiguous visualization of each transgene within individual cells. As expected, co-transfection of the two fluorescent payloads targeting the same LP (either *attP*-GT or *attP*-GA) resulted in cells expressing only one fluorophore, consistent with mutually exclusive integration. In contrast, co-transfection of *TurboGFP*-GT and *mScarlet-I*-GA donor plasmids yielded dual-labeled hiPSCs (**Figure 1g**). No integration was observed when the *TurboGFP*-GA donor was transfected into the parental *CLYBL*-bxb-v2 hiPSC line which lacks the GA allele, further supporting allele specificity and excluding detectable off-target integration (**Supplementary Figure 1i**). Finally, integrating the same fluorescent reporter into both LPs resulted in an ∼2-fold increase in fluorescence signal, suggesting that both *CLYBL* alleles support comparable levels of transgene expression (**Supplementary Figure 11**).

### Accelerating and simplifying the STRAIGHT-IN Dual protocol improves cell line generation

To streamline the generation of uniform, genetically modified hiPSCs, we optimized the STRAIGHT-IN integration process to improve efficiency, increase cell yield, and shorten the time before downstream experiments could be performed (**Figure 2a**). We hypothesized that the lower cytotoxicity and higher transgene expression typically observed with modRNA delivery compared to plasmid DNA in hiPSCs would improve integration outcomes. Indeed, transfection of Bxb1 modRNA resulted in an approximate 2-fold increase in integration efficiency (**Supplementary Figure 2a**).

**Figure 2.**
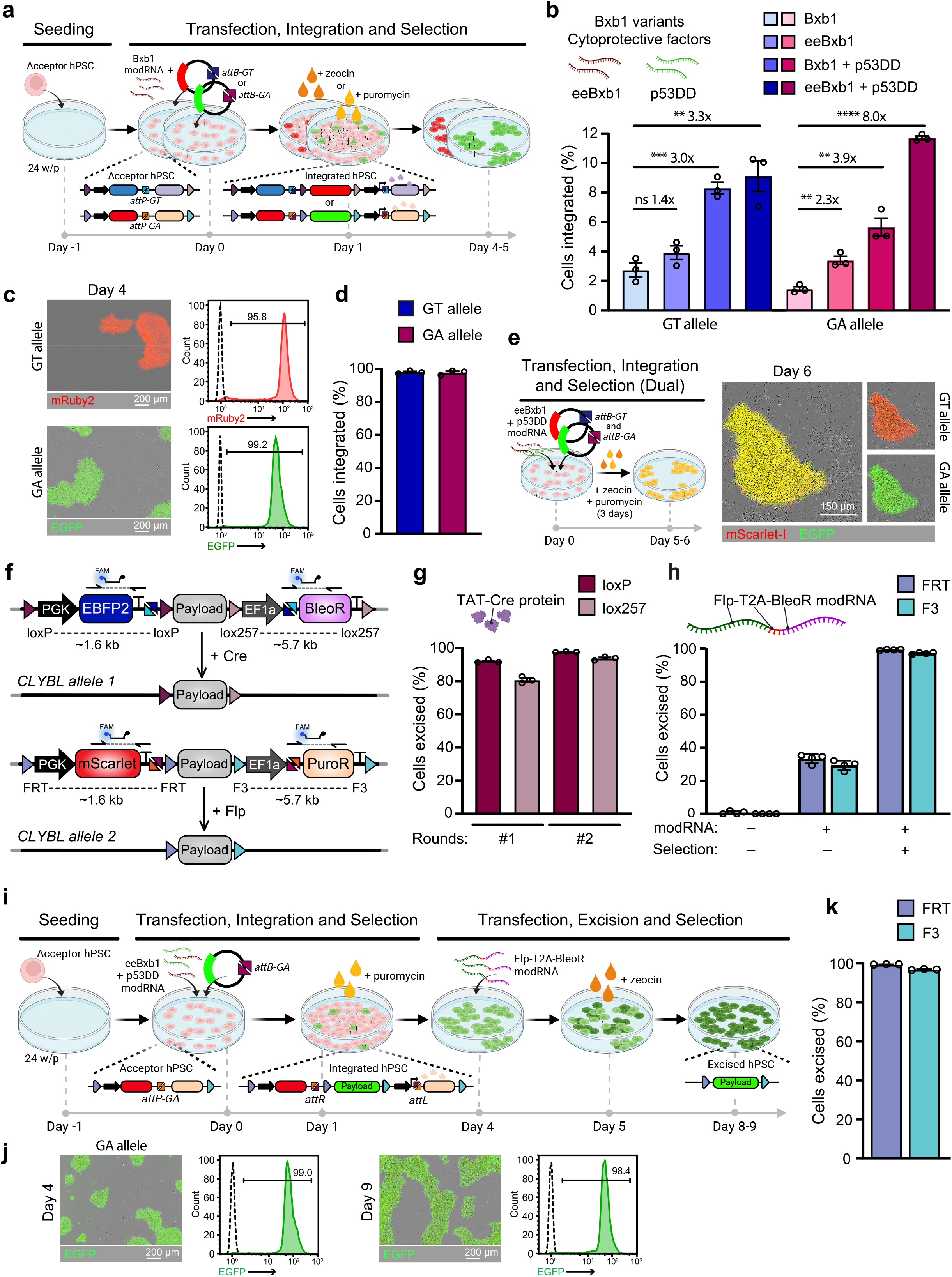
Optimized STRAIGHT-IN protocol improved integration and excision efficiencies while reducing timelines. (***a***) Schematic of the rapid integration procedure. (***b***) Mean integration efficiencies of GT and GA donor plasmids using modRNA combinations of Bxb1, eeBxb1 and p53DD prior to selection. N=3 independent transfections; error bars, ±SEM; not significant (ns), P>0.05; *, P≤0.05; **, P≤0.01; ***, P≤0.001; ****, P≤0.0001 (unpaired t test). (***c***) Overlay of fluorescence and phase contrast images (*left*) and flow cytometric analysis (*right*) showing reporter expression following rapid integration of GT or GA donor plasmids encoding *mRuby2* or *EGFP*, respectively. Dashed lines denote untransfected STRAIGHT-IN Dual acceptor hiPSCs. (***d***) Mean integration efficiencies of GT and GA donor plasmids following antibiotic selection. N=3 independent transfections; error bars, ±SEM. (***e***) Schematic of the dual payload integration and selection procedure (*left*), and overlay of fluorescence and phase contrast images (*right*) after co-delivery of GT and GA donor plasmids encoding *mScarlet-I* and *EGFP*, respectively. (***f***) Schematic for excising selection cassettes and plasmid backbones using Cre or Flp recombinases. Dashed lines indicate the sequences excised, and half arrows indicate primer sites for ddPCR analysis. (***g***) Mean percentages of hiPSCs with indicated flanking regions excised following 1 (#1) or 2 (#2) administrations of TAT-Cre protein, as determined by ddPCR. N=3 independent transfections; error bars, ±SEM. (***h***) Mean percentages of hiPSCs with indicated flanking regions excised following Flp-2TA-BleoR modRNA transfection, with (+) or without (-) zeocin selection, as determined by ddPCR. N=4 independent transfections; error bars, ±SEM. (***i***) Schematic of the complete rapid integration and excision workflow. (***j***) Representative fluorescence and phase contrast images, and flow cytometric analysis of hiPSCs transfected with a GA-*EGFP* donor plasmid on days 4 and 9 of the STRAIGHT-IN rapid integration and excision workflow. Dashed lines represent untransfected STRAIGHT-IN Dual acceptor hiPSCs. (***k***) Mean percentage of hiPSCs with indicated flanking regions excised after Flp-2TA-BleoR modRNA transfection and zeocin selection, as determined by ddPCR. N=3 independent transfections; error bars, ±SEM.

Next, we explored whether co-delivery of cytoprotective factors could improve post-transfection survival, as reported for CRISPR-based genome editing (Haideri et al., 2022, 2024). Co-transfection of *p53DD* modRNA increased the proportion of hiPSCs with an integrated cargo by 12.4-fold compared to a control *mGreenLantern* modRNA (**Supplementary Figure 2b**). We also evaluated four engineered Bxb1 variants with reported improvements in recombination activity (Hew et al., 2024; Pandey et al., 2025), namely evoBxb1 (V74A), eeBxb1 (V74A + E229K + V375I), Bxb1-I87L, and Bxb1-I87L + A369P + E434G. Among these, eeBxb1 delivered as modRNA, and combined with p53DD modRNA, significantly increased pre-selection integration efficiency from 2.75% to 9.12% at the GT allele, and from 1.46% to 11.7% at the GA-allele (**Figure 2b**). Other Bxb1 variants did not outperform wildtype Bxb1, although co-transfection of *p53DD* modRNA consistently improved integration efficiencies across all experiments (**Supplementary Figure 2c**).

To expedite enrichment using zeocin or puromycin, we utilized modRNA for faster Bxb1 expression compared to plasmid DNA, allowing selection to begin one day after transfection. This reduced the time required to obtain uniform, payload-carrying hiPSCs to less than one week (**Figure 2c,d)**. Notably, inclusion of *p53DD* modRNA enabled recovery of hiPSC colonies with both GT and GA integrations after only 3 days of antibiotic selection (**Figure 2e**, **Supplementary Figure 2d**). ddPCR confirmed single-copy insertions of the GT and GA donor plasmids, with no evidence of random integration (**Supplementary Figure 2d**).

After integration, Cre and Flp recombinases are used to excise auxiliary elements, leaving behind only the DNA payload and often improving integrated transgene expression (**Figure 2f**)(Blanch-Asensio et al., 2022). While plasmid-based recombinase expression can result in high excision rates with selection, it often results in few clones surviving and prolonged expansion times. To improve this, we explored plasmid-free strategies using either modRNA or recombinant protein delivery.

For the GT allele, delivery of TAT-Cre protein resulted in excision rates of 91.9% and 80.7% of the *loxP-* and *lox257*-flanked sequences, respectively. A second round of TAT-Cre increased this to 97.6% and 93.7% (**Figure 2g**). For the GA allele, where TAT-Flp protein is unavailable, we used *Flp* modRNA. After one transfection, 61.5% and 54.8% of the hiPSCs had excised the *FRT-* and *F3*-flanked sequences, respectively, increasing to 87.8% and 76.3% with a second transfection (**Supplementary Figure 2f**, *left*). Co-delivery of *TAT-Cre* and *Flp* modRNA resulted in 83.6% of hiPSCs excising all flanked auxiliary sequences after two transfection rounds (**Supplementary Figure 2f**, *right*).

We hypothesized that coupling excision to selection might further enhance efficiency, potentially enabling complete removal of auxiliary elements after a single transfection. To test this, we integrated a donor plasmid into the GA allele, conferring puromycin resistance but keeping the cells zeocin sensitive. Transfecting a modRNA encoding both Flp and BleoR (*Flp-2TA-BleoR*), followed by zeocin selection starting one day later, resulted in nearly complete excision of both upstream and downstream auxiliary elements within 3 days with efficiencies of 99.3% and 97.2%, respectively (**Figure 2h**). However, an equivalent strategy could not be replicated with Cre for donor plasmids integrated at the GT allele, as puromycin selection following *Cre-T2A-PuroR* or *Cre-IRES-PuroR* modRNA transfection failed to yield puromycin-resistant hiPSCs. Co-transfection of *Cre* and *PuroR* modRNAs resulted in limited excision enrichment, with efficiencies only increasing from 25.7% to 44.3% for *loxP-* and from 22.5% to 41.7% for *lox257*-flanked sequences (**Supplementary Figure 2g**).

Finally, to further streamline the protocol, we performed donor plasmid integration and auxiliary element excision in the same well without needing to passage the cells. The hiPSCs were co-transfected with *eeBxb1* and *p53DD* modRNA along with a GA donor plasmid encoding *EGFP*, followed by puromycin selection days 1-3 post-transfection. On day 4, y *Flp-T2A-BleoR* modRNA was transfected, and zeocin selection applied from days 5-8 (**Figure 2i**). Within 9 days, hiPSCs showed near uniform EGFP expression and excision of auxiliary elements (**Figure 2j,k**), demonstrating a rapid, one-well protocol for precise, near-scarless single-copy genome integration with >98% efficiency.

### Identifying promoter sequences supporting stable transgene expression in hiPSCs

Transgenes are essential tools for tracking cellular processes and modulating cell behavior and identity (Peterman et al., 2024). However, in hiPSCs, genome-integrated transgenes are susceptible to transcriptional silencing, particularly during prolonged culture or upon differentiation. Promoter choice plays a critical role in determining both expression strength and silencing susceptibility (**Figure 3a**)(Cabrera et al., 2022; Seczynska et al., 2022). Although the CMV early enhancer/chicken β-actin (CAG) and human elongation factor 1-alpha (hEF1a) promoters are commonly used for constitutive transgene expression across diverse cell types (Dou et al., 2021), their comparative activity in hiPSCs remains poorly defined.

**Figure 3.**
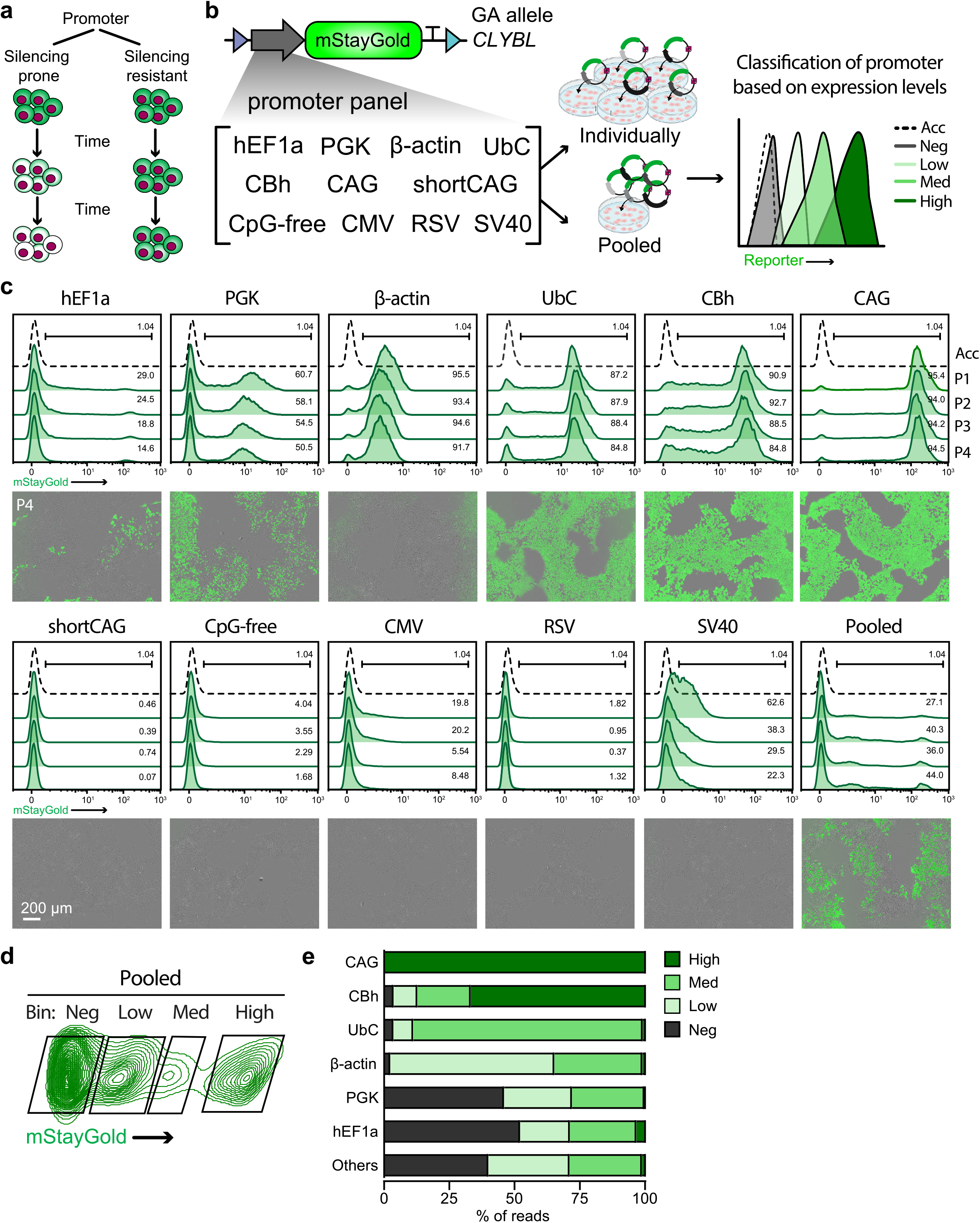
Evaluation of transgene silencing using STRAIGHT-IN. (***a***) Conceptual schematic illustrating transgene silencing over time following genomic integration. (***b***) Schematic of the panel of 11 different promoters, each driving expression of a *mStayGold* reporter integrated into the *CLYBL* locus. Donor plasmids were delivered either individually or as a pooled library to identify promoter sequences supporting high, medium and low transgene expression levels. (***c***) Representative flow cytometric analysis (*top*), and overlaid fluorescence and phase-contrast images (*bottom*) over 4 passages of *mStayGold* expression in hiPSCs following individual promoter integration. Dashed lines indicate untransfected STRAIGHT-IN Dual acceptor hiPSCs. (***d***) Flow cytometric analysis of the pooled integration approach, with the bulk population sorted into four *mStayGold* expression clusters: negative (neg), low, medium (med), and high. (***e***) Bar graph showing the distribution of sequencing reads for the indicated promoter sequences mapping to each of the four expression clusters.

To directly compare their activity, we generated a dual reporter hiPSC line in which divergently oriented CAG and hEF1a promoters drove expression of *mRuby2* and *mGreenLantern*, respectively. While all puromycin-resistant hiPSC colonies showed uniform mRuby2 expression, only a subset expressed mGreenLantern despite confirmed integration of the hEF1a-driven cassette, indicating transcriptional silencing. This effect became more pronounced with passaging (**Supplementary Figure 3a**).

To exclude reporter-specific effects and promoter interference, we replaced *mGreenLantern* with *mStayGold*, the brightest monomeric GFP reported to date (H. Zhang et al., 2024). Additionally, we integrated the two reporters into separate *CLYBL* alleles. Again, only the hEF1a-driven mStayGold was silenced (**Supplementary Figure 3b**). Swapping promoter-reporter pairs also confirmed this, with strong expression of CAG-driven mStayGold while hEF1a-driven mRuby2 was silenced (**Supplementary Figure 3c**).

We next screened a panel of 11 promoters to identify sequences capable of driving low, medium, or high levels of transgene expression. In addition to hEF1a and CAG, the panel included ubiquitous promoters (UbC, β-actin, PGK), viral (CMV, RSV, SV40) promoters, truncated CAG variants (CBh, shortCAG), and CpG-depleted hEF1a variant (CpG-free). We introduced the promoters either separately or in a multiplexed format to demonstrate the feasibility of integrating plasmid libraries using STRAIGHT-IN. This also enabled direct comparison of pooled versus individual outcomes (**Figure 3b**). While all promoters initially produced detectable mStayGold expression 24 h after transfection, their expression profiles diverged markedly following single-copy genomic integration at the *CLYBL* locus, indicating potential context-dependent differences between episomal and integrated transgene expression (**Supplementary Figure 4a,b**).

**Figure 4.**
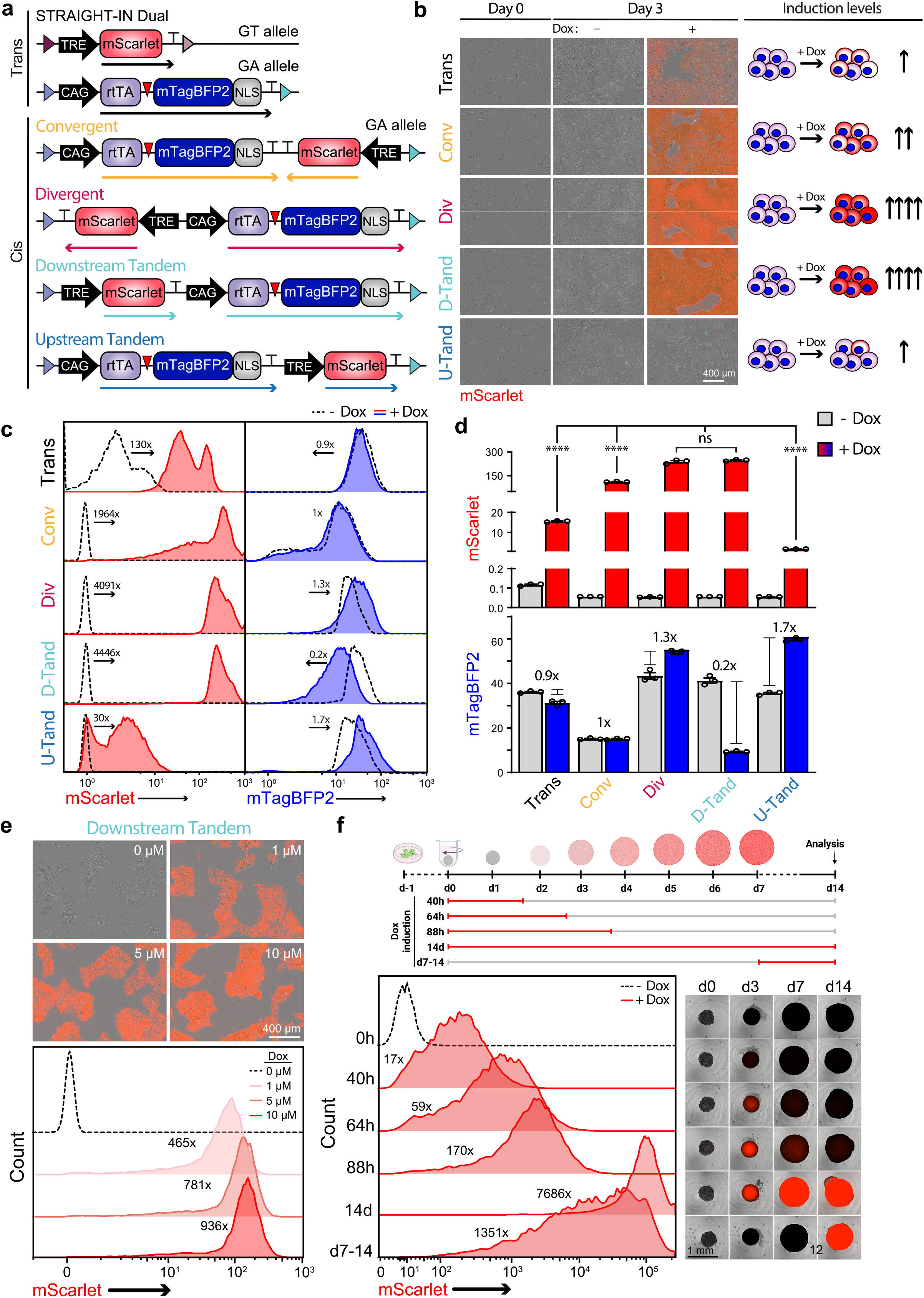
Gene syntax modulates performance of the Tet-On 3G system in hiPSCs. (***a***) Schematic overview of a trans design and the four possible all-in-one Tet-On 3G syntaxes, defined by the relative orientation and order of the constitutive (*rtTA*-T2A-*mTagBFP2*) and doxycycline-inducible (TRE- *mScarlet*) transcriptional units. Arrows indicate transcriptional direction. (***b***) Representative fluorescence/phase-contrast images (*left*) of hiPSCs carrying the integrated constructs shown in (**a**), cultured in the absence (-) or presence (+) of doxycycline for 3 days. The schematic (*right*) summarizes relative *mScarlet* expression across syntaxes. (***c***) Flow cytometric analysis of *mScarlet* and *mTagBFP2* expression in hiPSCs carrying each of the integrated constructs, in the absence (*dashed line*) or presence (*red*/*blue*) of doxycycline for 3 days. Values indicate fold change which is based on geometric mean fluorescence intensity (*G-mean*). (***d***) Quantification of G-mean values for *mScarlet* and *mTagBFP2* expression in the absence (*grey*) or presence (*red*/*blue*) of doxycycline for 3 days. N=3 biological replicates; mean ±SEM; ns, P>0.05; ****, P≤0.0001 (one-way ANOVA). (***e***) Dose–response of doxycycline-induced *mScarlet* expression in hiPSCs harbouring the downstream tandem construct. Representative fluorescence/phase-contrast images (*top*) and flow cytometric analysis with values indicating fold changes based on the G-mean (*bottom*). (***f***) Inducible *mScarlet* expression in cardioids carrying the downstream tandem construct. Schematic of the induction protocols (*top*). Flow cytometric analysis showing *mScarlet* expression in the absence (*dashed line*) or presence (*red*) of doxycycline for the indicated periods (*bottom left*). Fold change values (x) are based on the G- mean values relative to untreated cardioids. Representative fluorescence/phase-contrast images at different timepoints for the same indicated periods of doxycycline induction (*bottom right*).

Among the candidates tested, full-length CAG consistently supported the strongest expression of mStayGold, followed by its truncated derivative CBh. The UbC and β-actin promoter sequences produced intermediate and low expression levels respectively, suggesting their potential utility in applications requiring tighter control. In contrast, the remaining promoters, including hEF1a, CMV, and CpG-free, either failed to promote homogenous mStayGold expression or were rapidly silenced upon passaging (**Figure 3c**). Longitudinal monitoring over ten passages showed sustained mStayGold expression from CAG and UbC promoters, while with hEF1a it remained silenced in most cells (**Supplementary Figure 4c**).

We next evaluated promoter behavior in a pooled context. All 11 promoter constructs were detected in the bulk population by next generation sequencing (**Supplementary Figure 4d**). Cells were then binned into four mStayGold-based expression clusters (negative, low, medium, and high), and the specific promoter sequences in each sorted cluster were mapped (**Figure 3d,e** and **Supplementary Figure 4e**). The expression patterns closely matched those from individually integrated lines, with CAG and CBh enriched in the high expression cluster, UbC in the intermediate, β-actin in the low, and hEF1a, PGK and the remaining promoters predominantly appearing in the negative cluster.

To assess whether these results were locus-dependent, we repeated the analysis by integrating 6 of the promoter constructs (hEF1a, CAG, UbC, CBh, β-actin, PGK) into a hiPSC line containing a STRAIGHT-IN LP at the *AAVS1* locus (Blanch-Asensio et al., 2023) (**Supplementary Figure 5a**). As observed at the *CLYBL* locus, CAG drove strong and stable expression across multiple passages (**Supplementary Figure 5b**). Notably, UbC and CBh showed greater variability in silencing and levels between loci, indicating both the promoter and the locus influence the stability and strength of transgene expression.

**Figure 5.**
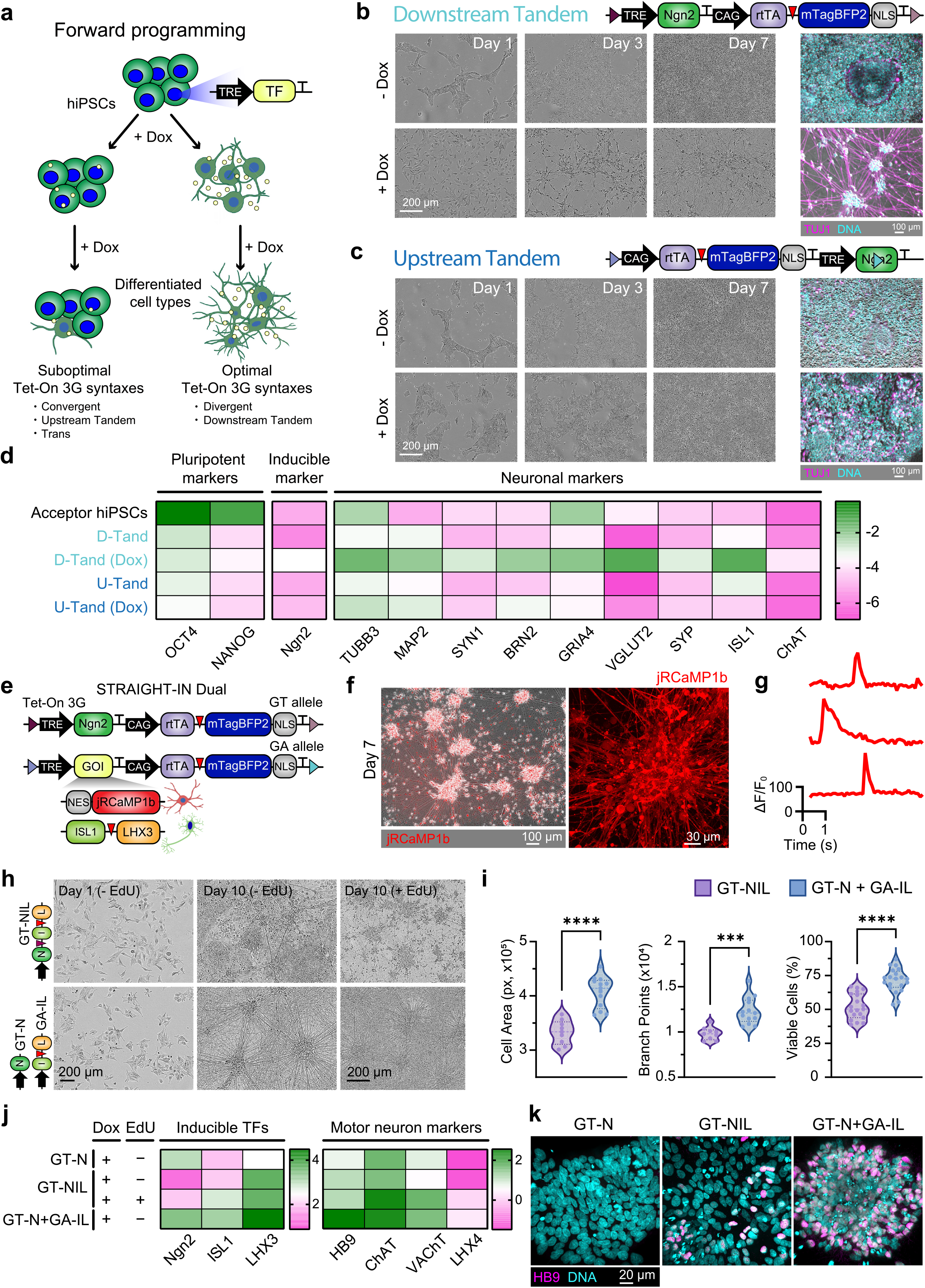
Doxycycline-inducible expression of multiple payloads using STRAIGHT-IN Dual. (***a***) Schematic of forward programming strategy to generate specific cell types by inducible overexpression of transcription factors. (***b***) Schematic of a downstream tandem all-in-one doxycycline-inducible *Ngn2* cassette (*top*). Phase contrast and immunofluorescence images (*TUJ1*, magenta; *DNA*, cyan) of cells at indicated days treated with (+) or without (-) doxycycline (*bottom*). (***c***) Schematic of an upstream tandem all-in-one doxycycline-inducible Ngn2 cassette (*top*), with corresponding images to those shown in **b** (*bottom*). (***d***) Expression analysis of pluripotency and neuronal marker genes in untransfected hiPSCs and cells containing inducible *Ngn2* cassettes from **(b)** or **(c)**, treated with (+) or without (-) doxycycline for 7 days. Values are normalized to *RPL37A* and log_10_-transformed. N=3 independent differentiations. (***e***) Schematic of STRAIGHT-IN Dual hiPSCs with the inducible *Ngn2* cassette from **(b)** in the GT allele, and a similar downstream tandem construct containing either a *jRCaMP1b* or a bicistronic *ISL1-LHX3* cassette in the GA allele. (***f***) Representative fluorescence/phase-contrast (*left*) and *jRCaMP1b* fluorescence (*right*) images of forward programmed iNs at day 7. (***g***) Representative *jRCaMP1b* fluorescence traces showing cytosolic Ca^2+^ transients from forward programmed iNs. (***h***) Phase contrast images of cells containing the indicated transcription factor cassettes for motor neuron induction at day 1 and 10 of treatment with doxycycline ± EdU for 48 h. (***i***) Violin plots showing quantified cell area (*left*), number of branch points (middle), and percentage of viable (*right*) iMNs obtained using either the single- or dual-cassette configurations indicated. (***j***) Expression analysis of inducible transcription factors and motor neuron markers in cells containing GT-N, GT- NIL, or GT-N + GA-IL constructs, cultured with doxycycline for 10 days. Values are normalized to *RPL37A* and shown relative to uninduced conditions (log_10_-transformed). N=3 independent differentiations. (***k***) Immunofluorescence images (*HB9*, magenta; *DNA*, cyan) of GT-N, GT-NIL and GT-N+GA-IL cells treated with doxycycline for 10 days.

These findings guided further optimization of the STRAIGHT-IN donor plasmids. Since CAG consistently supported strong and stable transgene expression and hEF1a was more prone to silencing, we designed the GT and GA donor plasmids to place the CAG promoter upstream of the transgene cloning site (**Supplementary Figure 6a**). Additionally, as the original GA donor plasmid relied on hEF1a to drive *PuroR* expression in the LP, we hypothesized that this might limit the recovery of puromycin-resistant colonies. To test this, we replaced hEF1a with CAG, UbC or CBh (**Supplementary Figure 6b**). Following puromycin selection, we observed 1.45x, 2.29x and 3.84x increases in the colony numbers respectively, indicating that poor hEF1a activity was impairing selection efficiency (**Supplementary Figure 6c**).

**Figure 6.**
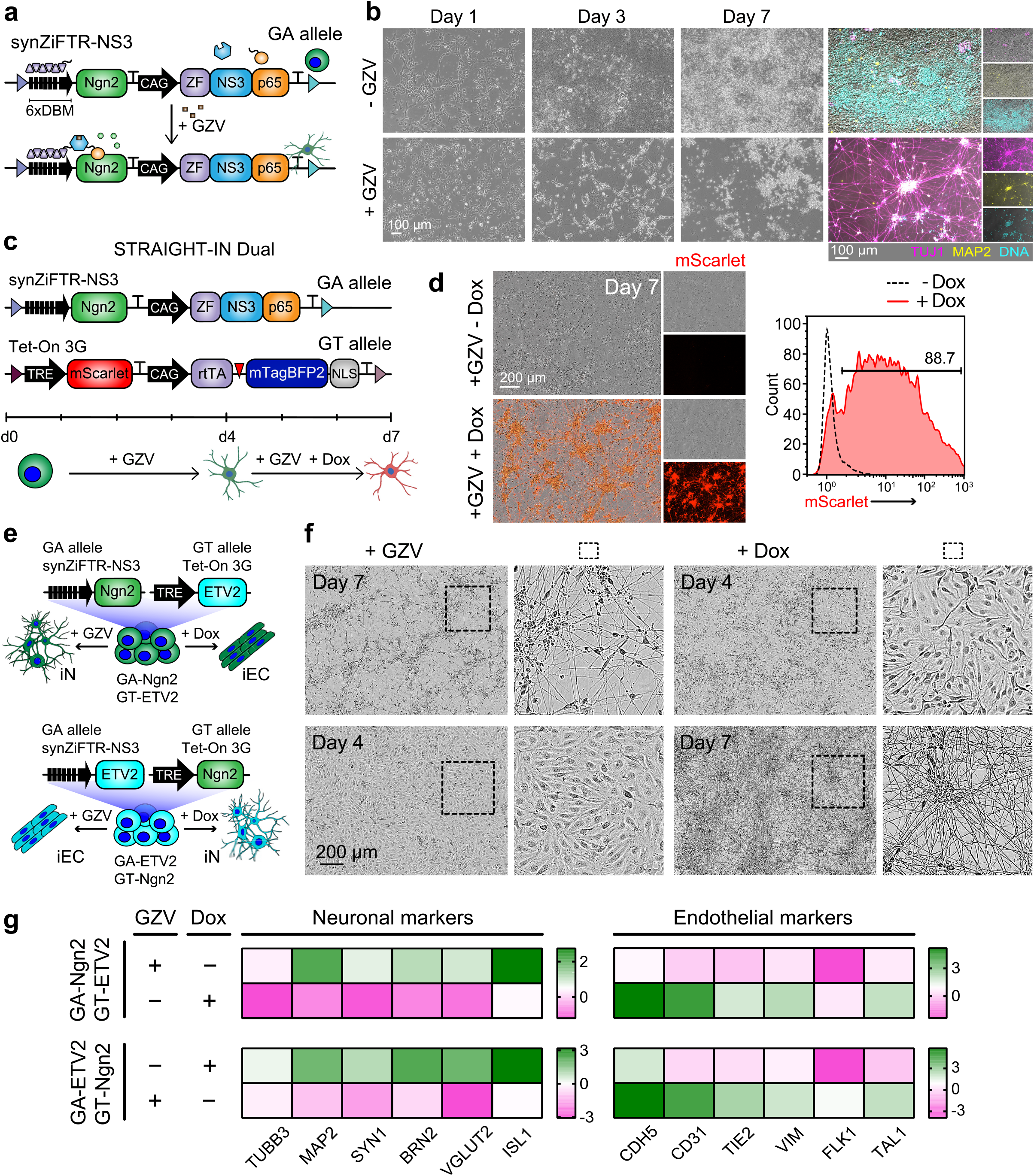
Dual-fate programming via orthogonal, inducible transcription factor expression using STRAIGHT-IN Dual. (***a***) Schematic of a downstream tandem synZiFTR-based *Ngn2* cassette integrated into the GA allele, enabling GZV-inducible expression. (***b***) Phase contrast and immunofluorescence images of cells at indicated time points, cultured with (+) or without (-) GZV (*TUJ1*, magenta; *MAP2*, yellow; *DNA*, cyan). (***c***) Schematic of STRAIGHT-IN Dual hiPSCs with a doxycycline-inducible *mScarlet* cassette in the GT allele and the GZV-inducible *Ngn2* cassette in the GA allele, enabling sequential and combinatorial induction. (***d***) Representative fluorescence and phase contrast images (*left*), and flow cytometric analysis (*right*) of GZV- induced iNs cultured with (*red line*) or without (*dashed line*) doxycycline. (***e***) Schematic of dual-fate STRAIGHT-IN Dual lines with all-in-one downstream tandem inducible *Ngn2* or *ETV2* cassettes regulated by synZiFTR (GA allele) or Tet-On 3G (GT allele) systems. (***f***) Representative phase contrast images of hiPSC directed into iNs or iECs in the presence of either GZV or doxycycline for the indicated days. (***g***) Gene expression analysis of neuronal and endothelial markers from the cells in (**f**). Values are normalized to *RPL37A* and shown relative to uninduced condition (log_10_-transformed). N=3 independent differentiations.

### Gene syntax influences induction efficiency in the Tet-On 3G system in hiPSCs

Beyond promoter selection, the relative arrangement of transcriptional units, often referred to as gene syntax, can influence the expression of adjacent genes (Engreitz et al., 2016; Johnstone & Galloway, 2022; Patel et al., 2023). While placing different elements of genetic circuits, such as inducible systems, at separate genomic loci may prevent suboptimal interactions between genes, integration at separate sites also requires integrating additional genetic cargo. To systematically compare how different designs of multi-component genetic circuits perform, we used STRAIGHT-IN Dual to elucidate key design principles, focusing on the Tet-On 3G system and examining trade-offs between co-localized and dual locus integration.

We first constructed a *trans* design of the Tet-On 3G system, with the transcriptional units integrated into the separate *CLYBL* alleles (**Figure 4a**). The inducible gene (*mScarlet*) was driven by a tetracycline response element (TRE) containing seven tetracycline operator (*TetO*) repeats. Having determined the CAG promoter resists silencing in hiPSCs, we used this promoter to drive expression of the transactivator gene *rtTA* in a bicistronic cassette which also expressed a nuclear-localized blue fluorescent reporter (*mTagBFP2-NLS)* as a proxy readout for rtTA levels.

We also generated *cis* designs of the Tet-On 3G system, where both the constitutive (*rtTA-T2A-mTagBFP2-NLS*) and inducible (*mScarlet*) transcriptional units were integrated into the same allele. This allowed us to investigate how gene syntax, specifically the relative orientation and order of the transcriptional units, affects induction. We created four possible syntaxes: convergent, divergent, downstream tandem and upstream tandem (**Figure 4a**). As the auxiliary sequences remaining in the GA allele could confound potential interactions between the constitutive and inducible genes, we examined mTagBFP2 and mScarlet expression both before (**Supplementary Figure 7**) and after excision of the sequences (**Figure 4**).

In the *trans* configuration, induced mScarlet expression was weak, whereas several *cis* designs showed robust induction (**Figure 4b**), suggesting that spatial proximity between the constitutive and inducible genes improves expression in the Tet-On 3G system. Notably, *mTagBFP2* expression levels remained unchanged in the *trans* configuration, indicating that resource burden was not a limiting factor in induction (**Figure 4c,d**).

Prior to induction, we observed clear bimodality in mTagBFP2 expression for the convergent syntax, which was most pronounced in unexcised cells (**Supplementary Figure 7c**), and aligned with biophysical models of transcription (Johnstone & Galloway, 2022). In contrast, the other syntaxes exhibited unimodal mTagBFP2 expression, which remained stable upon induction with 1 µM doxycycline (**Supplementary Figure 7c**). Interestingly, in pre-excised cells, induction resulted in slight repression of mTagBFP2 across all syntaxes (**Supplementary Figure 7d**). However, after excision, syntax-specific differences in mTagBFP2 expression emerged (**Figure 4c,d**). Most notably, the downstream tandem syntax exhibited an ∼5-fold reduction in mTagBFP2 expression, while the convergent syntax showed no significant change. The divergent and upstream tandem syntaxes showed increased mTagBFP2 expression, in line with biophysical model predictions (**Figure 4c,d**).

Upon induction with 1 µM doxycycline, striking syntax-specific differences also emerged in mScarlet expression. Although these differences became more pronounced after excision (**Figure 4b-d**), similar trends were observed before excision (**Supplementary Figure 7e,f**). The divergent and downstream tandem syntaxes resulted in strong mScarlet induction, while the convergent and upstream tandem syntaxes showed poorer induction, characterized by weak, bimodal, or broad expression distribution. Interestingly, excision significantly improved induction for the convergent syntax, possibly due to the increased expression of mTagBFP2 and therefore rtTA. However, excision did not improve induction for the upstream tandem syntax (**Figure 4c,d**).

Given the robust mScarlet induction observed in the downstream tandem *cis* design, we further explored its utility in 3D stem cell-derived cardiac organoids (cardioids), which provide a more complex cellular environment for disease modeling and developmental studies. We found that mScarlet expression could be modestly adjusted by varying doxycycline concentration (**Figure 4e**). Additionally, altering the duration of doxycycline treatment modulated mScarlet expression, with longer inductions leading to higher expression (**Figure 4f**). Even in developed cardioids, robust mScarlet expression could be achieved when adding doxycycline from day 7, demonstrating the potential to dynamically modulate transgene expression in complex 3D systems.

### Dual payload inducible system enables forward programming of hiPSCs into iNs and iMNs

Forward programming of hiPSCs into specific cell types is typically achieved by overexpressing lineage-defining transcription factors. However, suboptimal gene syntax may impair transcription factor expression, inhibiting cell fate conversion (**Figure 5a**). Given the striking differences we observed between the two tandem orientations (**Figure 4d**), we investigated whether the gene syntax influenced forward programming outcomes when overexpressing *Ngn2*, a pioneer transcription factor for neurons (Y. Zhang et al., 2013). To evaluate this, we constructed Tet-On 3G all-in-one systems for regulating Ngn2 expression using either the downstream or upstream tandem syntaxes.

Induction of *Ngn2* from the downstream tandem syntax led to the rapid generation of TUJ1^+^ MAP2^+^ hiPSC-derived iNs within 7 days (**Figure 5b** and **Supplementary Figure 8a**). In contrast, the upstream tandem configuration produced very few iNs (**Figure 5c**), consistent with its weaker expression profile observed earlier (**Figure 4c,d**). qPCR analysis confirmed reduced induction of *Ngn2* and other neuronal genes in the upstream tandem syntax, while pluripotency-associated genes were downregulated in the downstream tandem orientation after doxycycline induction (**Figure 5d**).

To demonstrate the utility of STRAIGHT-IN Dual for co-expressing multiple transgenes, we integrated *Ngn2* and a genetically-encoded calcium indicator (*jRCaMP1b*) into the separate *CLYBL* alleles using two TRE-controlled donor plasmids (**Figure 5e**). With doxycycline as a single inducer, we achieved rapid and robust forward programming of hiPSCs into iNs while simultaneously recording calcium signals (**Figure 5f,g** and **Supplementary Figure 8b**).

We next investigated whether distributing the transcription factors required for motor neuron (iMN) specification (*Ngn2, ISL1* and *LHX3*) across both LPs would improve the homogeneity of forward programming outcomes. Specifically, we compared a dual-cassette configuration in which *Ngn2* and the *ISL1/LHX3* cassette were integrated separately (GT-Ngn2 + GA-ISL1- LHX3; GT-N + GA-IL; **Figure 5e**) to a single-cassette configuration where all three factors were encoded in tandem within a single LP (GT- Ngn2-ISL1-LHX3; GT-NIL). The single-cassette GT-NIL configuration led to a high proportion of proliferative, non-neuronal cells that rapidly overtook the culture (**Figure 5h** and **Supplementary Figure 8c**). In contrast, the dual-cassette strategy resulted in improved neuronal differentiation, yielding a purer population of iMNs (**Figure 5h** and **Supplementary Figure 8c**). In the single-allele approach, the non-neuronal cells could be selectively eliminated by treatment with the antimitotic reagent EdU, a step not required for the dual-cassette design (**Figure 5h** and **Supplementary Figure 8d**).

After 10 days of differentiation, Calcein AM staining and image quantification revealed that cells derived from the dual-cassette configuration were significantly larger, had increased neurite branching, and longer dendrites (**Figure 5i** and **Supplementary Figure 9**). Live-cell imaging with Hoechst and propidium iodide showed increased cell death in EdU-treated single-cassette cultures, consistent with the higher proportion of non-neuronal, EdU-sensitive cells (**Figure 5i** and **Supplementary Figure 9c,d**). Furthermore, qPCR confirmed that expression of *Ngn2, ISL1*, and *LHX3* was lower in the single-cassette configuration, while the dual-cassette cultures showed robust upregulation of motor neuron markers, including HB9, ChAT and VaChT, compared to *Ngn2* alone (**Figure 5j** and **Supplementary Figure 9e**). HB9 protein expression was not detected in the absence of *ISL1*, and *LHX3* (GT-N cultures), but was detectable in the single-cassette and most strongly expressed in the dual-cassette cultures (**Figure 5k**).

For rapid cloning and integration of genes of interest (GOIs) into the STRAIGHT-IN LPs, we developed a set of downstream tandem and divergent Tet-On 3G donor plasmids (**Supplementary Figure 10a**). These vectors include a constitutive nuclear BFP reporter for visualization and a LacZ cassette to facilitate blue/white screening of bacterial colonies that contained the cloned GOIs. After assembly, most white colonies (>85%) carried the correct cargo (**Supplementary Figure 10b,c**). This streamlined cloning and screening approach allowed for the rapid generation of donor plasmids with diverse inducible cargoes that could then be integrated using the rapid protocol into the STRAIGHT-IN alleles. These hiPSC lines, which were established in under one week, also uniformly expressed the nuclear BFP reporter (**Supplementary Figure 10d,e**).

### Rapid generation of hiPSC-iECs and promoter activity profiling in differentiated lineages using STRAIGHT-IN Dual

To further demonstrate the versatility and efficiency of our all-in-one Tet-On 3G downstream-tandem vector system for generating inducible hiPSC lines, we cloned and overexpressed *ETV2*, a pioneer transcription factor that directs endothelial lineage specification (**Supplementary Figure 11a**). Upon doxycycline induction, cells rapidly acquired an endothelial-like morphology and, by day 4, expressed key markers of endothelial identity, including CD31, ZO1, and CD144 (**Supplementary Figure 11b-d**).

To evaluate promoter activity following lineage commitment, we integrated the previously characterized panel of 11 promoters into the GA allele of hiPSC lines harboring doxycycline-inducible *Ngn2* or *ETV2* in the GT allele. This enabled side-by-side comparison of promoter activity in iNs and iECs, as well as hiPSCs (**Supplementary Figure 12)**.

Promoter behavior in iNs and iECs largely mirrored that observed in undifferentiated hiPSCs. As in the pluripotent state, the CAG promoter drove strong and uniform mStayGold expression in both iNs and iECs (**Supplementary Figure 12b,c**). Similarly, the UbC and β-actin promoters also remained active, producing moderate and low levels of reporter expression, respectively. In contrast, the remaining promoters, including hEF1a, PGK and CBh, either failed to produce detectable reporter signals or exhibited transgene silencing, consistent with observations in undifferentiated hiPSCs.

### Orthogonal inducible systems support dual-fate programming of genetically uniform hiPSCs

To achieve independent control over two separate cargoes within a single hiPSC line, we introduced a second, orthogonal inducible system alongside the Tet-On 3G system. We selected a synZiFTR system which couples ZF10 as a DNA-binding domain to a grazoprevir (GZV)- responsive NS3 module and a p65 transcriptional activator (H.-S. Li et al., 2022). An all-in-one synZiFTR expression cassette was constructed in the downstream tandem configuration and integrated into the GA allele (**Supplementary Figure 13a,b**). Induction of a *mGreenLantern* reporter revealed toxicity at GZV concentrations above 250 nM, with 125 nM selected as the optimal concentration for robust transgene induction with minimal cytotoxicity (**Supplementary Figure 13c**). At this concentration, synZiFTR-mediated Ngn2 expression efficiently generated TUJ1^+^/MAP2^+^ iNs within 7 days (**Figure 6a**,b).

To demonstrate independent dual gene regulation, we combined both inducible systems using STRAIGHT-IN Dual. *Ngn2* was integrated into the GA allele under synZiFTR control, while mScarlet was targeted to the GT allele under Tet-On 3G regulation (**Figure 6c**). GZV was maintained in the culture media throughout the 7- day differentiation procedure, with doxycycline added from day 3. By day 7, most iNs expressed mScarlet (**Figure 6d** and **Supplementary Figure 13d**).

To exploit this modularity, we established dual-fate hiPSC lines capable of differentiating into distinct lineages based on inducer choice. In one configuration, *Ngn2* expression was controlled by the synZiFTR system and *ETV2* by Tet-On 3G, while in another, this arrangement was reversed (**Figure 6e**). Exposure to GZV or doxycycline resulted in rapid and efficient differentiation into either iNs or iECs (**Figure 6f**). Lineage-specific commitment was supported by qPCR analysis and flow cytometry, with GZV-induced iECs expressing CD31 and CD144 (**Figure 6g** and **Supplementary Figure 11e**). These findings highlight the versatility of STRAIGHT-IN Dual for independently controlling multiple transcriptional programs in a single, genetically uniform hiPSC line. This capability provides a flexible framework for implementing orthogonal gene circuits and generating mixed or patterned cell populations for advanced applications.

## Discussion

In this study, we present STRAIGHT-IN Dual, a platform enabling rapid, efficient, and allele-specific integration of two independent DNA payloads into the *CLYBL* locus in hiPSCs. Following protocol optimization, genetically modified hiPSC lines could be generated in less than one week with nearly 100% efficiency. For the GA allele, both integration and excision procedures were achievable within nine days without requiring cell passaging.

STRAIGHT-IN Dual supports simultaneous integration of two payloads with orthogonal precision, exploiting the highly specific GT and GA Bxb1 recombination sequences (Jusiak et al., 2019). By using eeBxb1, a hyperactive Bxb1 mutant developed via directed evolution (Pandey et al., 2025), we achieved significantly higher integration efficiencies compared to wild-type Bxb1. Notably, this improvement was more pronounced at the GA allele. This may be due to the attP/attB-GA sequence being included in some of the circuits used during the directed evolution of Bxb1 to eeBxb1 (Pandey et al., 2025), potentially biasing the enzyme toward enhanced recombination at these sites. However, the precise mechanism by which the central dinucleotide sequence influences recombination efficiency remains unclear and requires further investigation.

Since Bxb1 catalyzes recombination through DNA double-strand breaks (Xu et al., 2013), we also investigated whether co-delivering modRNA encoding a dominant-negative truncated p53 (p53DD) to dampen the p53-mediated DNA damage response would enhance cell survival after recombination (Haideri et al., 2024; M. Li et al., 2022; Rosenstein et al., 2024). Combining eeBxb1 and p53DD resulted in approximately a 10-fold increase in integration efficiency.

The platform also supports near-scarless genomic modifications by excising auxiliary elements, such as reporters, selection markers, and plasmid backbones. Consistent with prior findings (Blanch-Asensio et al., 2024), removal of these auxiliary sequences reduced transgene silencing in both undifferentiated and differentiated hiPSCs. Excision also allows reuse for further genomic modifications. STRAIGHT-IN Dual facilitated near-complete excision efficiency from both alleles using TAT-Cre and *Flp-T2A- BleoR* modRNA, with the entire procedure for the GA allele achieved without cell passaging. This streamlined approach simplifies the generation of multiple modified hiPSC lines, either individually or pooled, facilitating integration of plasmid libraries and high-throughput combinatorial studies at single-copy resolution. These optimized processes also make STRAIGHT-IN Dual compatible with automated pipelines, making it ideal for large-scale functional screens and high-throughput studies.

Importantly, dual integration of the same payload into both alleles effectively doubled transgene expression, indicating that both LPs perform comparably. We recently demonstrated that such uniform biallelic integration is critical for high-resolution genome folding profiling and enables precise measurements of transcriptional activation effects on chromatin architecture using Region Capture Micro-C (Johnstone et al., 2025).

Our work also highlights current limitations of standard molecular tools used in hiPSCs. For example, the commonly used hEF1a promoter was rapidly silenced, corroborating earlier studies (Bertero et al., 2016). To identify promoter sequences that resist silencing, we screened a library of 11 promoters integrated either individually or as a pool into STRAIGHT-IN Dual hiPSCs. Next-generation sequencing of pooled integrations confirmed complete library representation, demonstrating the suitability of STRAIGHT-IN for scalable, high-throughput characterization of diverse DNA elements.

While transient assays initially showed detectable transgene expression from all promoters, over half underwent significant silencing following genomic integration. Viral promoters (CMV, RSV, SV40), a truncated CAG promoter variant (shortCAG), and a CpG-depleted short hEF1a promoter (CpG-free) were silenced rapidly, while PGK and hEF1a promoters showed progressive silencing over time. In contrast, CAG, CBh, UbC, and β-actin promoters maintained stable reporter expression at the *CLYBL* locus over extended passaging in hiPSCs, although only CAG maintained long-term activity at the *AAVS1* locus. When assessed in forward programmed neurons and ECs, most promoter behaviors mirrored that in undifferentiated hiPSCs. Notably, CAG, UbC, and β-actin promoters remained active, supporting their utility for stable gene expression in cell types derived from different germ layers.

The superior stability of the full-length CAG promoter likely arises from its synthetic architecture, which integrates several regulatory elements that collectively confer resistance to transcriptional silencing. CAG combines the cytomegalovirus (CMV) immediate-early enhancer, the chicken β-actin promoter with its first intron, and a rabbit β-globin splice acceptor (Niwa et al., 1991). These elements contain numerous transcription factor binding sites, can recruit chromatin remodelers and prevent DNA methylation (Brown et al., 2014). In contrast, truncated variants such as CBh, shortCAG, and CMV alone, lack at least the β-globin splice acceptor, and showed reduced resistance to silencing. Nonetheless, the large size and high GC content of CAG presents practical cloning challenges, highlighting the need to systematically develop and evaluate alternative synthetic promoters that retain the robust, stable expression characteristics of CAG but offer improved ease of use. STRAIGHT-IN Dual offers not only rapid, locus-controlled screening of promoter sequence libraries but also the possibility to evaluate other regulatory elements, such as insulators or ubiquitous chromatin opening elements, that have been proposed to mitigate transgene silencing in hiPSCs and their differentiated derivatives (Uenaka et al., 2025; Yanagi et al., 2025; Guichardaz et al., 2024).

We also demonstrated the utility of STRAIGHT-IN Dual for systematically analyzing gene syntax, identifying divergent and downstream tandem orientations as optimal configurations for induction with the all-in-one Tet-On 3G system. Most studies have favored these two orientations in mammalian cells (Jillette et al., 2019; Kelkar et al., 2020; Ng et al., 2021; Otomo et al., 2023; Randolph et al., 2017), with convergent and upstream tandem syntaxes typically performing poorly unless extensively optimized (Haenebalcke et al., 2013; Shin et al., 2024).

Based on the clear differences in expression observed between the two tandem syntaxes, we evaluated both for forward programming of hiPSCs into iNs via inducible *Ngn2* expression. We selected the downstream tandem syntax over the divergent orientation due to its higher transgene induction levels prior to excising the flanking auxiliary elements, thereby obviating the need to perform this step and allowing the initiation of overexpression studies within one week. Functional iNs were only obtained when using the downstream tandem syntax, confirming its superior performance. This syntax also proved effective for *ETV2* overexpression, enabling efficient differentiation of hiPSCs into iECs within four days. While similar outcomes have been reported previously, these studies relied on hyperactive transposase-based delivery systems that randomly inserted >50 copies of the construct throughout the genome and potentially led to unintended genomic disruption (Rieck et al., 2024; Ding et al., 2025). In contrast, we demonstrate that a single, site-specific integration of *ETV2* is sufficient to forward program the entire hiPSC population into iECs and suggests that high transgene copy numbers are not a prerequisite for rapid and uniform cell fate specification when expression is tightly controlled.

We found that distributing the transcription factors required for motor neuron programming across two separate alleles improved differentiation efficiency compared to encoding all three factors in a single multi-cistronic cassette (Wang et al., 2025). This design may improve expression of individual factors by avoiding position-dependent effects common to 2A-linked polycistronic constructs (Liu et al., 2017). STRAIGHT-IN Dual therefore enables systematic screening of transcription factor combinations for lineage specification, which previously was only feasible with pooled viral strategies that offer limited control over transcription factor stoichiometry (Joung et al., 2023; Ng et al., 2021).

Moreover, STRAIGHT-IN Dual allows combinatorial or sequential transgene expression using independent inducible systems, such as Tet-On 3G and synZiFTRs. This enabled precise temporal control of both reporters and lineage-specifying transcription factors. Using this dual-inducible framework, we generated dual-fate programmable hiPSC lines that could be directed toward either neuronal or endothelial lineages, depending on inducer choice. Previous multi-fate approaches have relied on mixing or printing genetically distinct cell populations (Skylar-Scott et al., 2022). In contrast, our strategy enables spatial patterning of distinct lineages from a single cell line using small molecules alone, without the need for complex bioprinting. The platform also opens possibilities to combine cell fate programming with independent expression of biosensors or inducible modeling of sporadic or acquired mutations in differentiated cells. Furthermore, these inducible DNA payloads can be expressed in complex in vitro systems, such as organoids, organs-on-chips, and 3D tissue constructs, expanding the possibilities for cell-based modelling and engineered tissue design.

Several limitations of the current study should be acknowledged. First, all engineered hiPSC lines were derived from a single parental clone. Although beneficial for reducing variability, this genetic uniformity limits the immediate generalizability of our findings. Further validation using hiPSC lines from diverse genetic backgrounds and donors is necessary. Second, most payloads were integrated into the *CLYBL* locus, with limited comparisons at *AAVS1*. Therefore, locus-specific effects cannot be excluded, and transgene behavior could differ across genomic sites. Third, the transgene sequence also influences transgene silencing (Karbassi et al., 2024). Here, promoter activity was primarily assessed using fluorescent reporter genes. Further testing of promoter activity for a broader range of relevant payloads is warranted in future studies.

In summary, STRAIGHT-IN Dual enables the rapid generation of complex, precisely engineered hiPSC lines with dual, orthogonal genetic control. This versatile platform supports scalable screening, combinatorial transgene interrogation, and precise control of cell fate, and provides a powerful toolkit for biomedical and synthetic biology research.

## Acknowledgements

We thank Niels Geijsen and Peng Shang for providing TAT-Cre protein, the LUMC flow cytometry facility for sorting the cells, and the Laboratory for Diagnostic Genome Analysis (LUMC) for karyotyping. Some schematics in figures were created with BioRender.com. This work was supported by the Netherlands Organ-on-Chip Initiative, a Gravitation project of the Nederlandse Organisatie voor Wetenschappelijk Onderzoek (NWO), funded by the Ministry of Education, Culture, and Science of the government of the Netherlands (grant no. 024.003.001); the NWO-funded LymphChip project (grant no. NWAORC 2019 1292.19.019) as part of the NWA research program ‘‘Research on Routes by Consortia (ORC)’’; and a ZonMw PSIDER consortium grant (grant no. 10250022120002; GREAT). It also forms part of the project “Innovative Stem Cell Technology Infrastructure for Human Organ and Disease Models”, funded by the NWO Large-Scale Research Infrastructure program (grant no. 184.036.006). The Novo Nordisk Foundation Center for Stem Cell Medicine (reNEW) is supported by a Novo Nordisk Foundation grant (NNF21CC0073729). Additional support was provided by the National Institute of General Medical Sciences of the National Institutes of Health (award no. R35-GM143033 to K.E.G), the National Science Foundation under the NSF-CAREER award (grant no. 2339986), and the Institute for Collaborative Biotechnologies. This study also received funding from European Research Council (ERC) under the European Union’s Horizon 2000 research and innovation program (grant no.101042634).

## Author contributions

A.B-A developed and optimized the protocols described in this manuscript, designed and performed the experiments, analyzed data and wrote the manuscript. D.S.P developed and optimized some of the protocols described in this manuscript, designed and performed some of the experiments, contributed to drafting the manuscript and revised it for important intellectual content. S.C, B.B.J, M.B and N.B.W, designed and performed some of the experiments, and contributed to drafting the manuscript. V.O contributed intellectual content. A.A analyzed data, contributed to drafting the manuscript and revised it for important intellectual content. C.L.M acquired some of the funding and revised the manuscript for important intellectual content. K.E.G and R.P.D. supervised the study, acquired some of the funding and revised the manuscript for important intellectual content. All authors approved the final manuscript.

## Notes

### Competing Interest Statement

The authors have declared no competing interest.

### Summary of Updates

Manuscript revised to include new results, updated in Figures 3, 5 and 6. Former Figure 3 has been moved to the Supplementary information as Sup Figures 3 and 6. Sup Figures 1, 6 and 8 contain new results. Sup Figures 4, 5, 9, 11, 12, 14 and 15 are new.

