## Supplemental file for "STRAIGHT-IN Dual: a platform for dual, single-copy integrations of DNA payloads and gene circuits into human induced pluripotent stem cell"

### Supplementary Figure 1

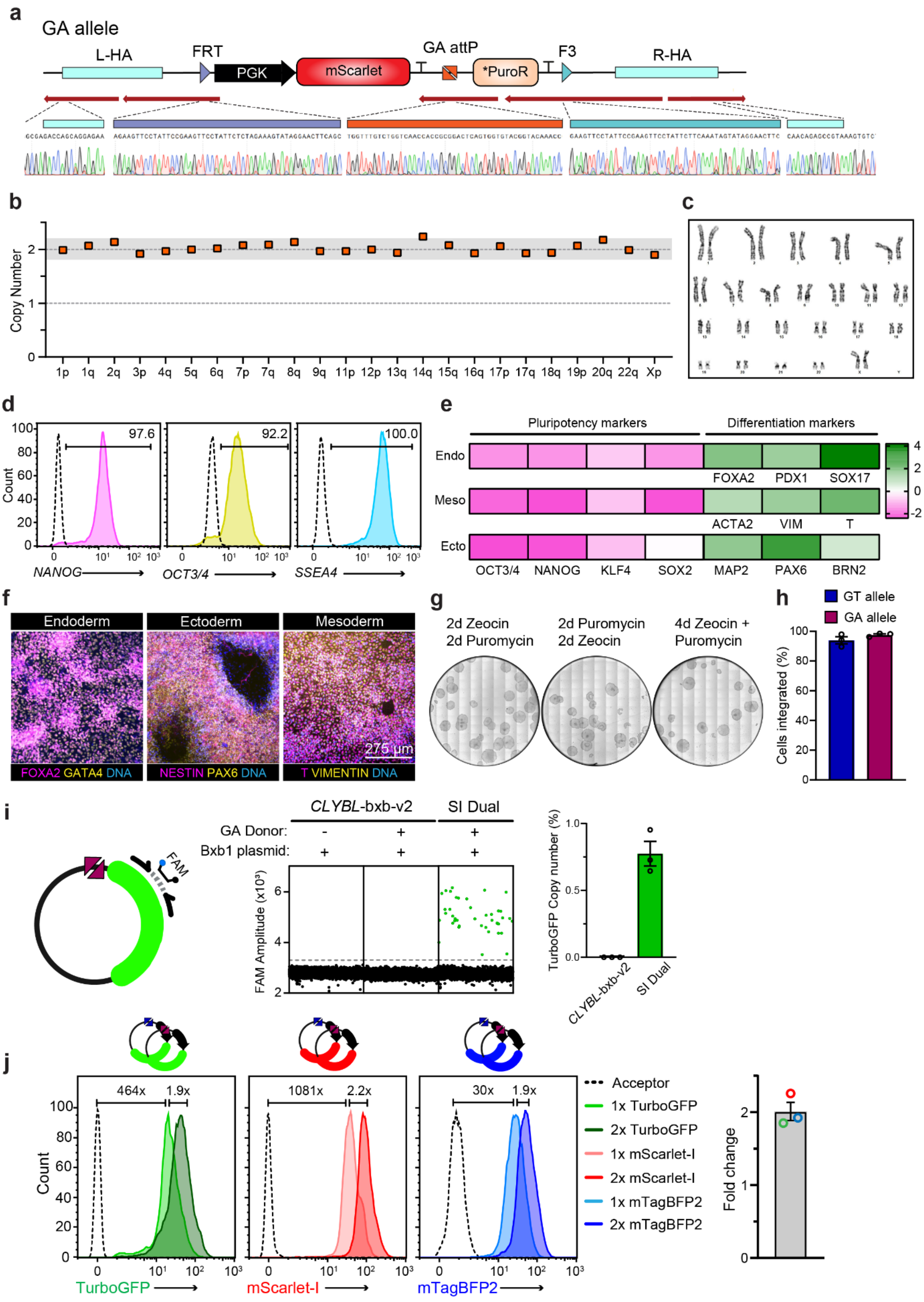

#### Supplementary Figure 1. Characterization of the STRAIGHT-IN Dual hiPSC line

**(a)** Sanger sequencing confirming correct targeting of the GA allele at the *CLYBL* locus, including sequences of the FRT, F3 and attP-GA sites. Red arrows indicate alignment of the sequencing chromatograms with the reference sequence. L-HA, left homology arm; R-HA, right homology arm.

**(b)** ddPCR analysis of 24 genomic loci covering >90% of the most common genetic abnormalities in hiPSCs. The STRAIGHT-IN Dual line showed a normal copy number (2) at all loci screened.

**(c)** G-banding karyogram confirming the STRAIGHT-IN Dual hiPSC line has a normal karyotype.

**(d)** Flow cytometric analysis of pluripotency-associated markers *NANOG*, *OCT3/4* and *SSEA-4* in the STRAIGHT-IN Dual hiPSC line. Dashed lines indicate unstained hiPSCs.

**(e)** Gene expression analysis of pluripotency-associated and lineage-specific markers following differentiation into the three germ layers. Values are normalized to *RPL37A* and presented relative to undifferentiated hiPSCs (log10-transformed). N=1 independent differentiation.

**(f)** Representative immunofluorescence images showing expression of germ layer markers: *FOXA2* and *GATA4* (endoderm), *NESTIN* and *PAX6* (ectoderm), and *T* and *VIMENTIN* (mesoderm) in differentiated STRAIGHT-IN Dual hiPSCs.

**(g)** Alkaline phosphatase staining of hiPSCs after sequential or simultaneous antibiotic selection to enrich for cells with both GT and GA donor plasmid integrations.

**(h)** Mean integration efficiencies of GT and GA donor plasmids co-delivered and selected simultaneously. N=3 independent transfections; error bars, ±SEM.

**(i)** Schematic of GA donor plasmid containing *TurboGFP* reporter (*left*) and representative ddPCR dot plots showing GA donor plasmid integration in the STRAIGHT-IN Dual line (*middle*). Green dots represent droplets containing the indicated sequence on the donor plasmid. Mean percentages of hiPSCs containing *TurboGFP* sequence (*right*). N=3 independent transfections; error bars, ±SEM.

**(j)** Flow cytometric analysis of STRAIGHT-IN Dual hiPSCs with one (1x) or two (2x) integrated fluorescent reporter payloads, as indicated. Dashed line indicates untransfected STRAIGHT-IN Dual acceptor hiPSCs. Fold change is the G-mean ratio. Right: mean fold change between hiPSCs containing 1 versus 2 integrated reporters. Error bars, ±SEM.

### Supplementary Figure 2

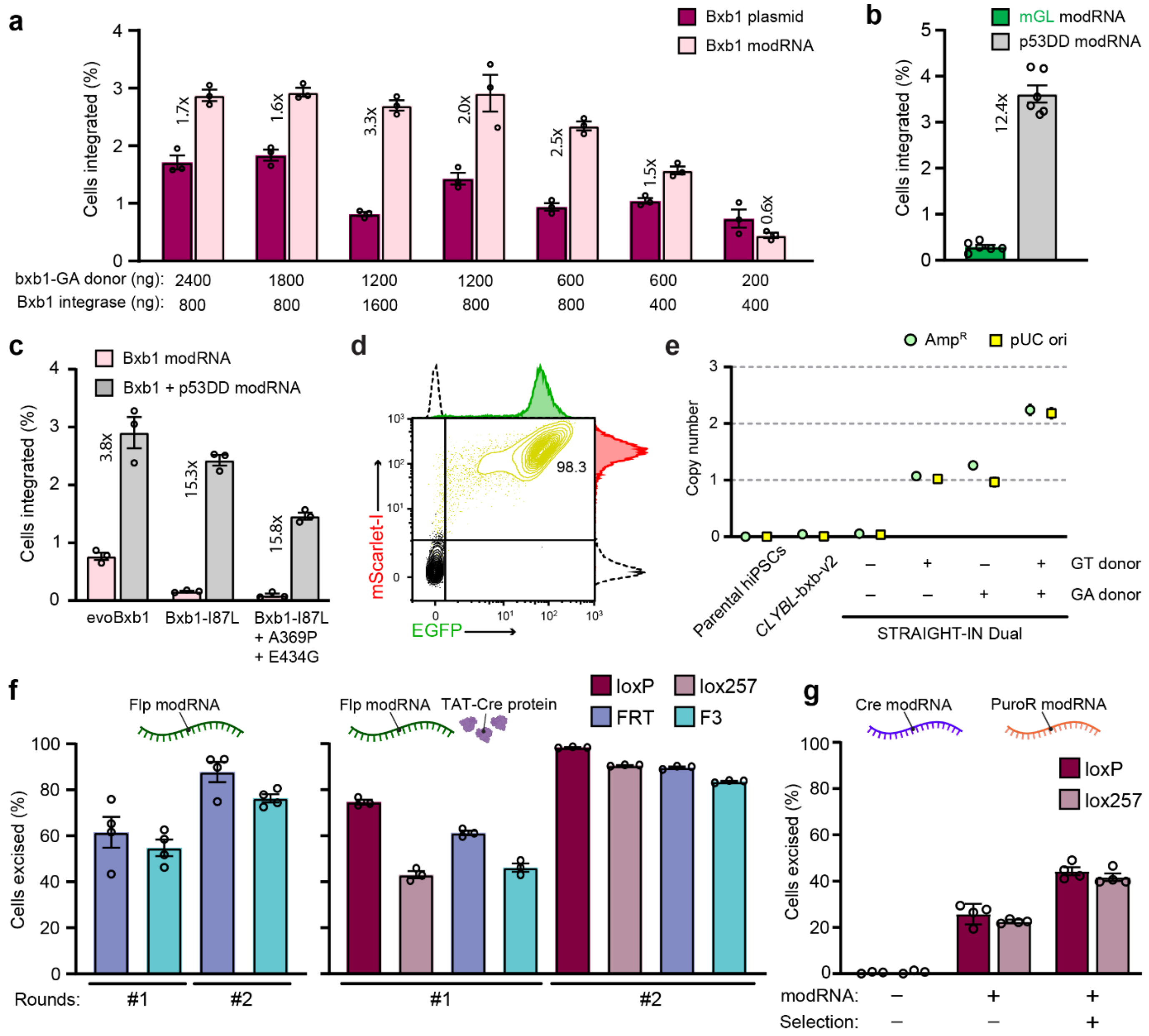

#### Supplementary Figure 2. Optimization of integration and excision processes in STRAIGHT-IN Dual hiPSCs

**(a)** Mean integration efficiencies (pre-selection) of a GA donor plasmid co-transfected with either Bxb1 expression plasmid or Bxb1 modRNA, with varying amounts of input. N=3 independent transfections; error bars, ±SEM.

**(b)** Mean integration efficiencies (pre-selection) of a GA donor plasmid co-transfected with Bxb1 expression plasmid and either mGreenLantern (mGL) or p53DD modRNA. N=6 independent transfections; error bars, ±SEM.

**(c)** Mean integration efficiencies (pre-selection) of a GA donor plasmid co-transfected with the indicated Bxb1 modRNA, with or without p53DD modRNA. N=3 independent transfections; error bars, ±SEM.

**(d)** Flow cytometric analysis of STRAIGHT-IN Dual hiPSCs 6 days after co-transfection of GT and GA donor plasmids encoding mScarlet-I and EGFP, respectively, along with Bxb1 and p53DD modRNAs, and following 3 days of antibiotic selection. Dashed line indicates untransfected hiPSCs.

**(e)** ddPCR analysis of AmpR and pUC ori backbone copy number in STRAIGHT-IN Dual hiPSCs following integration and selection with GT and/or GA donor plasmids. Error bars indicate Poisson 95% CI.

**(f)** Mean percentages of hiPSCs with excision of the indicated flanking regions following Flp modRNA alone (*left*), or TAT-Cre protein and Flp modRNA co-transfection (*right*), as determined by ddPCR. Recombinases were delivered once (#1) or twice (#2). N=3 independent transfections; error bars ±SEM.

**(g)** Mean percentages of excision of indicated flanking regions following co-transfection of Cre and PuroR modRNAs, with (+) or without (-) 24 h puromycin selection, as determined by ddPCR. N=4 independent transfections; error bars ±SEM.

#### Supplementary Figure 3

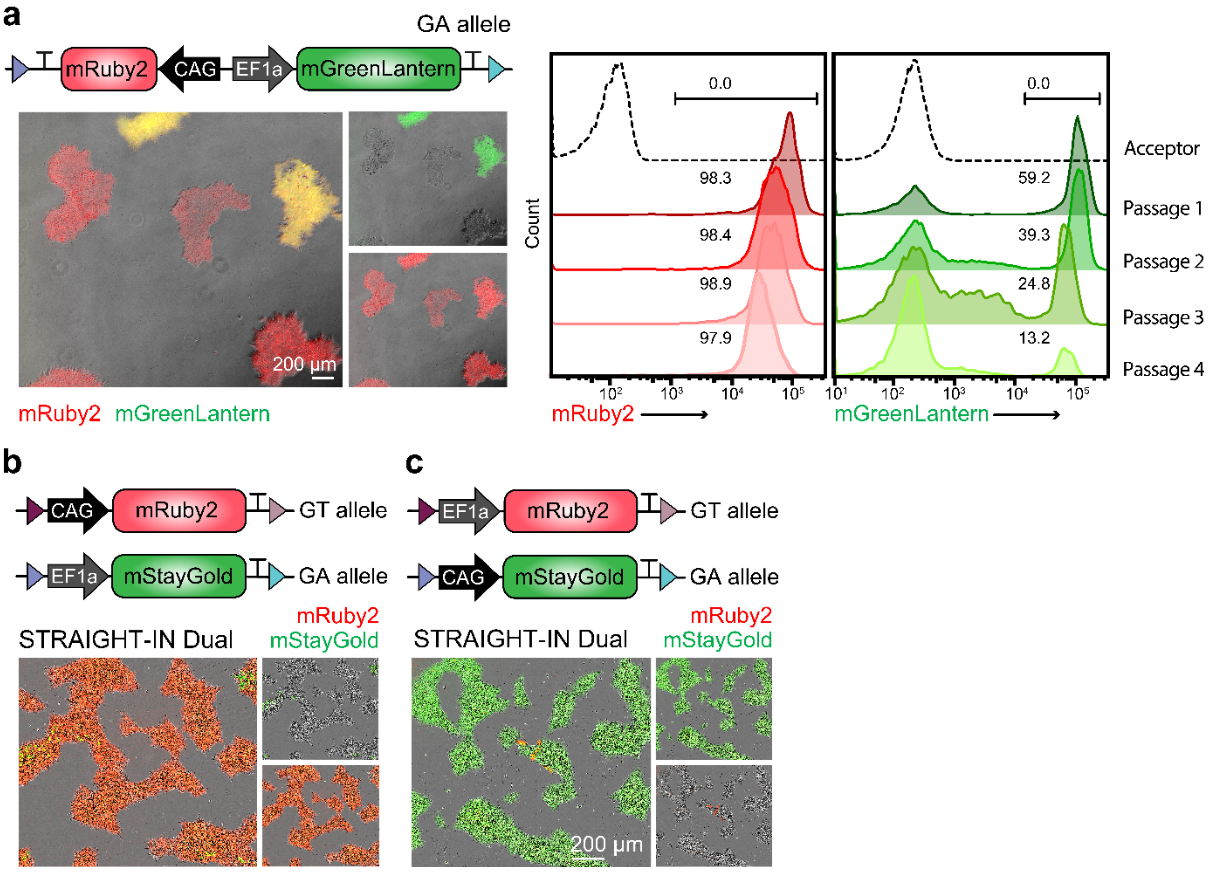


**Supplementary Figure 3. Transgenes driven by the hEF1a promoter are silenced in hiPSCs over time**

**(a)** Representative overlaid fluorescence and phase contrast images (*left*), and flow cytometric analysis over 4 passages (*right*), of hiPSCs expressing *mRuby2* and *mGreenLantern* reporters in cis, driven by the CAG and hEF1a promoters respectively, and integrated at the GA allele. Dashed lines represent untransfected hiPSCs.

**(b)** Overlaid fluorescence and phase contrast images (*left*), and flow cytometric analysis (*right*), of hiPSCs expressing mRuby2 and mStayGold reporters in trans, driven by the CAG and hEF1a promoters respectively, and integrated in the GT and GA alleles. Dashed lines represent untransfected hiPSCs.

**(c)** Overlaid fluorescence and phase contrast images of hiPSCs expressing mRuby2 and mStayGold reporters in trans, driven by the hEF1a and CAG promoters respectively, and integrated in the GT and GA alleles.

Supplementary Figure 4

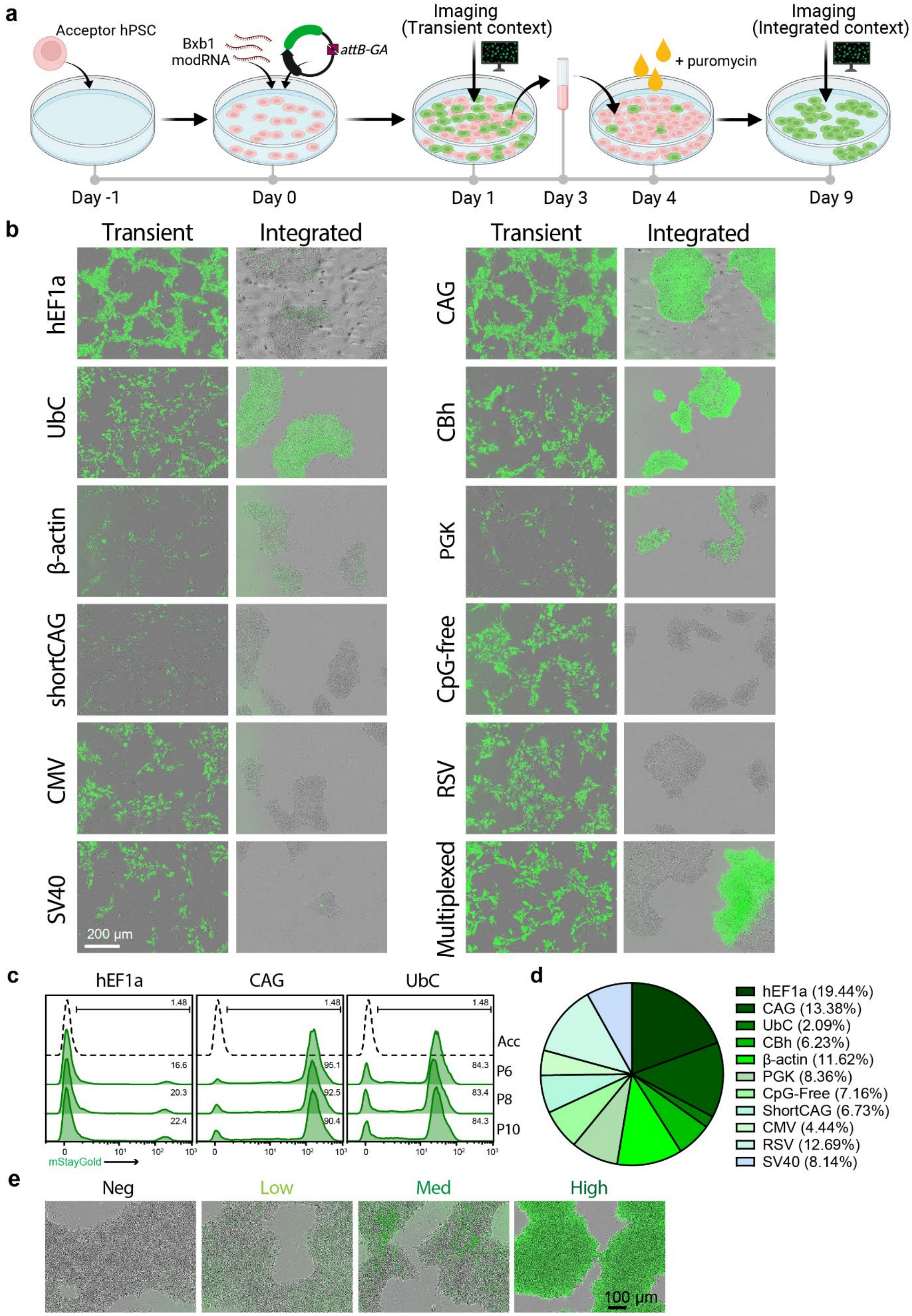

#### Supplementary Figure 4. Comparison of transient and integrated promoter-driven transgene expression in hiPSCs

**(a)** Schematic illustrating the experimental design. STRAIGHT-IN Dual hiPSCs were transfected with Bxb1 modRNA and GA donor plasmids encoding *mStayGold* under the control of diverse promoter sequences. Fluorescence was assessed 24 h post-transfection (transient expression) and again following puromycin selection and replating (genomic integrated expression).

**(b)** Representative fluorescence/phase-contrast images of hiPSCs expressing mStayGold driven under control of the indicated promoters, comparing transient and integrated expression.

**(c)** Flow cytometric analysis of mStayGold expression over 10 passages in hiPSCs with integrated constructs driven by hEF1a, CAG or UbC promoters.

**(d)** Pie chart of bulk population showing the percentage of sequencing reads corresponding to each of the 11 integrated promoter constructs in the pooled bulk population.

**(e)** Representative fluorescence/phase-contrast images of sorted bulk populations based on mStayGold expression levels (negative (neg), low, medium (med), and high), corresponding to the clusters shown in Figure 3d.

#### Supplementary Figure 5

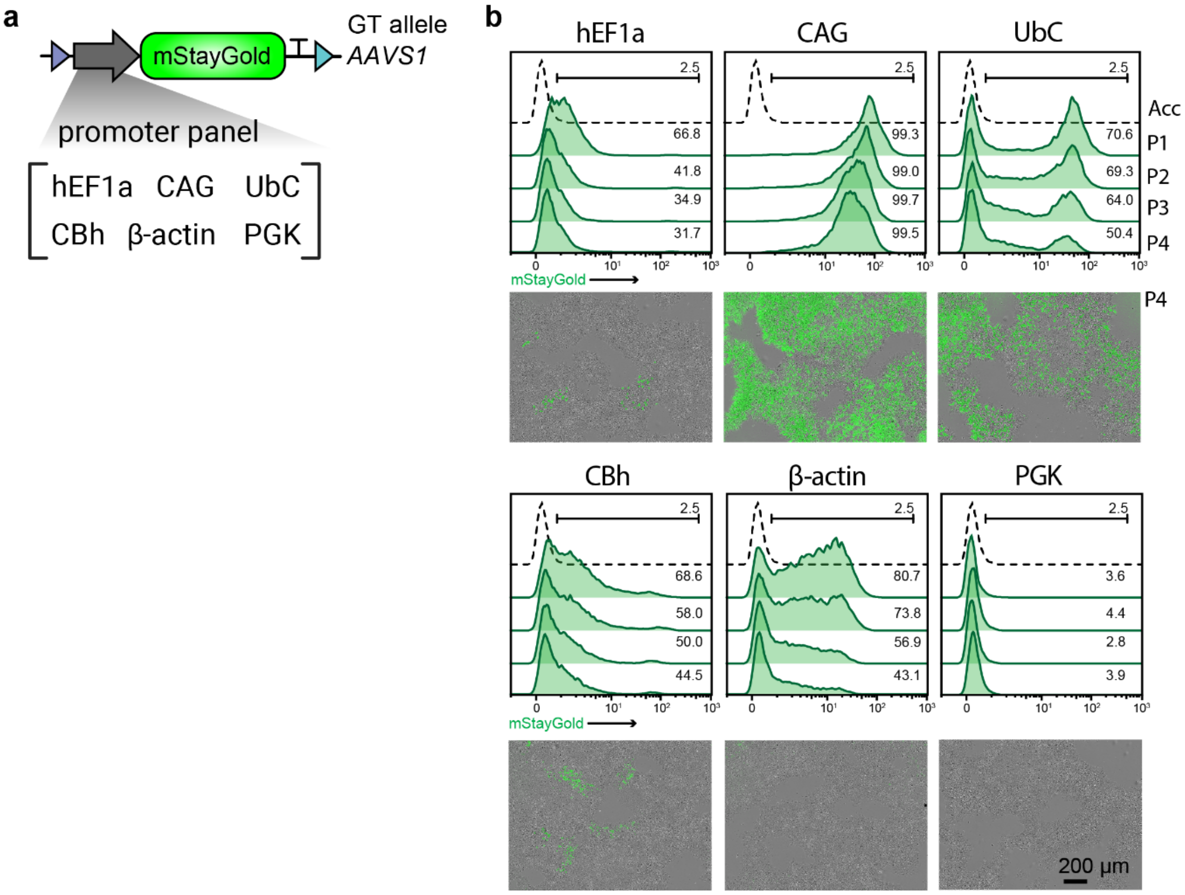


**Supplementary Figure 5. Characterization of promoter activity at the *AAVS1* locus**

(**a**) Schematic of the promoter panel, each driving expression of a *mStayGold* reporter and integrated at the *AAVS1* locus in STRAIGHT-IN *AAVS1* v2 acceptor hiPSCs.

(**b**) Flow cytometric analysis over 4 passages (*top*), and representative fluorescence/phase-contrast images at passage 4 (*bottom*), of hiPSCs expressing mStayGold from the different promoters indicated and integrated at the *AAVS1* locus, prior to excision of the auxiliary sequences. Dashed lines represent untransfected hiPSCs.

#### Supplementary Figure 6

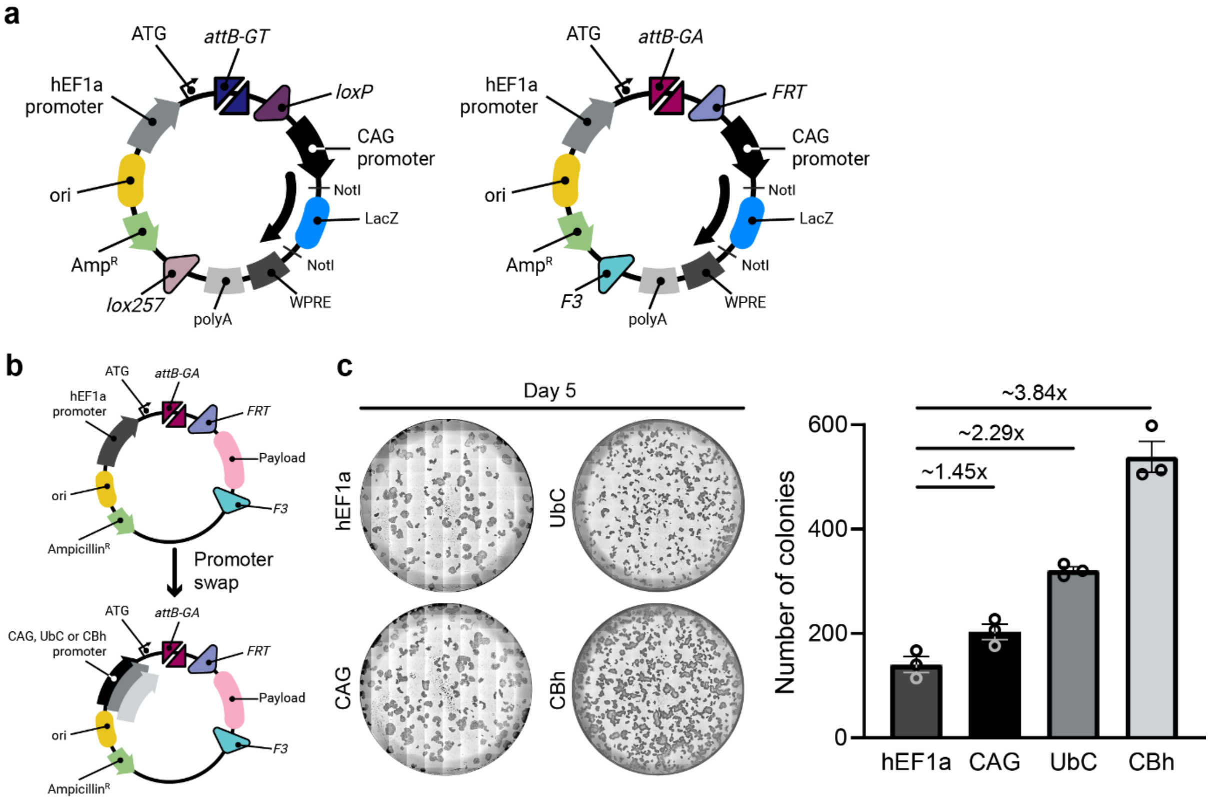


**Supplementary Figure 6. Optimization of STRAIGHT-IN donor plasmid backbone for improved transgene expression and integration efficiency**

**(a)** Schematic of the GT and GA donor plasmids containing a CAG promoter, WPRE and bGH polyadenylation signal for improved constitutive gene of interest (GOI) expression. The *LacZ* cassette is replaced with the GOI via NotI digestion followed by isothermal assembly or T4 ligation. White colonies on IPTG/X-gal agar plates identify clones containing the GOI.

**(b)** Schematic of GA donor plasmids in which the hEF1a promoter was substituted for CAG, UbC or CBh promoter sequences to assess the impact on integration efficiency.

**(c)** Representative phase contrast images (*left*) showing puromycin-resistant hiPSC colonies following donor plasmid integration, and quantification of the mean colony number per well (*right*). N=3 independent transfections; error bars, ±SEM.

#### Supplementary Figure 7

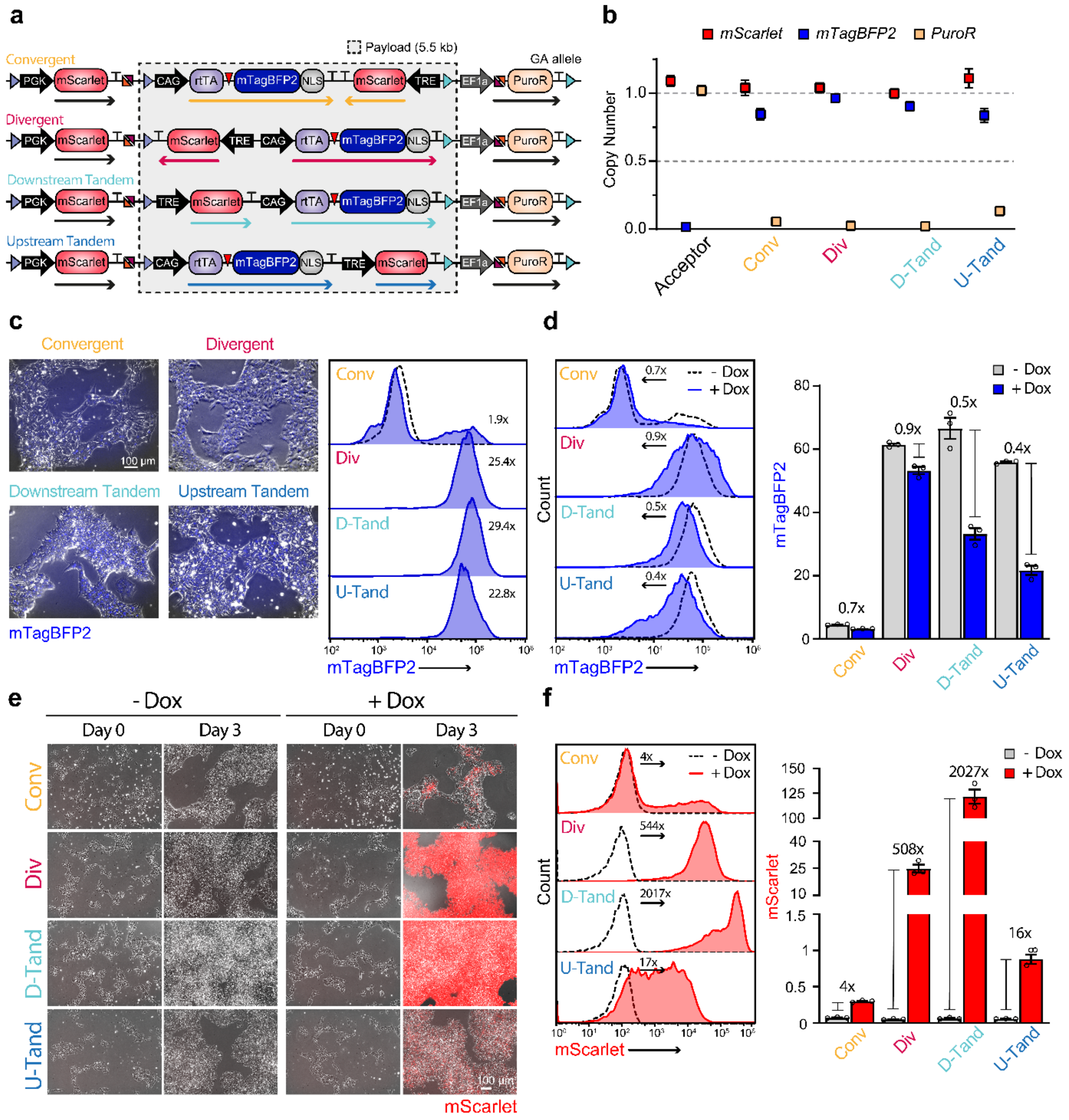

#### **Supplementary Figure 7. Evaluation of gene syntax configurations in an all-in-one Tet-On 3G inducible system**

**(a)** Schematic of the four cis-configured Tet-On 3G constructs, showing the orientation and order of the inducible (TRE-mScarlet) and constitutive (CAG-rtTA-T2A-mTagBFP2) cassettes, along with the upstream and downstream auxiliary elements. The grey-shaded region indicates the genomic sequence retained following Flp-mediated excision.

**(b)** ddPCR analysis confirming single-copy integration of *mScarlet* and *mTagBFP2*, and the absence of *PuroR* following excision. Error bars indicate Poisson 95% CI.

**(c)** Representative fluorescence/phase-contrast images (*left*) and flow cytometric analysis (*right*) of hiPSCs constitutively expressing mTagBFP2 from the constructs in (**a**) prior to excision. Fold change was calculated by comparing the G-mean relative to untransfected hiPSCs (dashed line), with values shown in plots.

**(d)** Flow cytometric analysis of mTagBFP2 expression in the absence (dashed line) or presence (blue) of doxycycline for 3 days (*left*), and quantification of G-mean fluorescence values (*right*). Fold change values (x) are calculated relative to untreated samples. N=3 biological replicates; mean ±SEM.

**(e)** Representative fluorescence/phase-contrast images of hiPSCs expressing inducible mScarlet from the constructs in (**a**) in the absence or presence of doxycycline for 3 days.

**(f)** Flow cytometric analysis of mScarlet expression in the absence (dashed line) or presence (red) of doxycycline for 3 days (*left*), and quantification of G-mean fluorescence values (*right*). Fold change values (x) are calculated relative to untreated samples. N=3 biological replicates; mean ±SEM.

Supplementary Figure 8

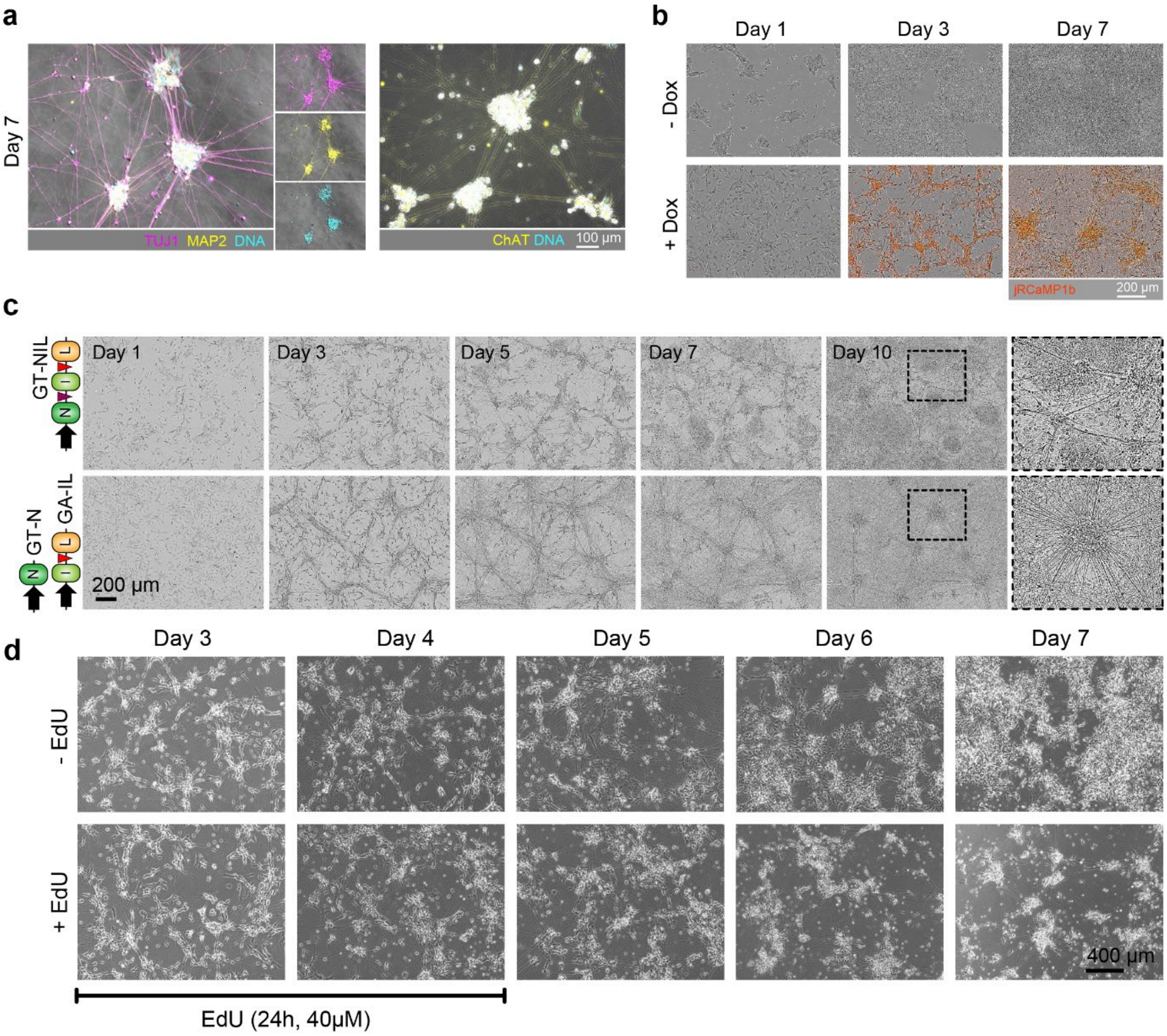


**Supplementary Figure 8. Forward programming of hiPSCs to neurons or motor neurons using the Tet-On 3G system with downstream tandem orientation**

**(a)** Overlaid immunofluorescence and phase contrast images showing *TUJ1* (magenta), *MAP2* or *ChAT* (yellow), and DNA (cyan) in cells cultured with doxycycline for 7 days.

**(b)** Representative fluorescence/phase-contrast images of cells cultured in the presence or absence of doxycycline for the indicated time points.

**(c)** Phase contrast images of cells over 10 days of culture in the presence of doxycycline.

**(d)** Phase contrast images of cells at different time points cultured with or without EdU administered for 48 hours between days 3–5.

#### Supplementary Figure 9

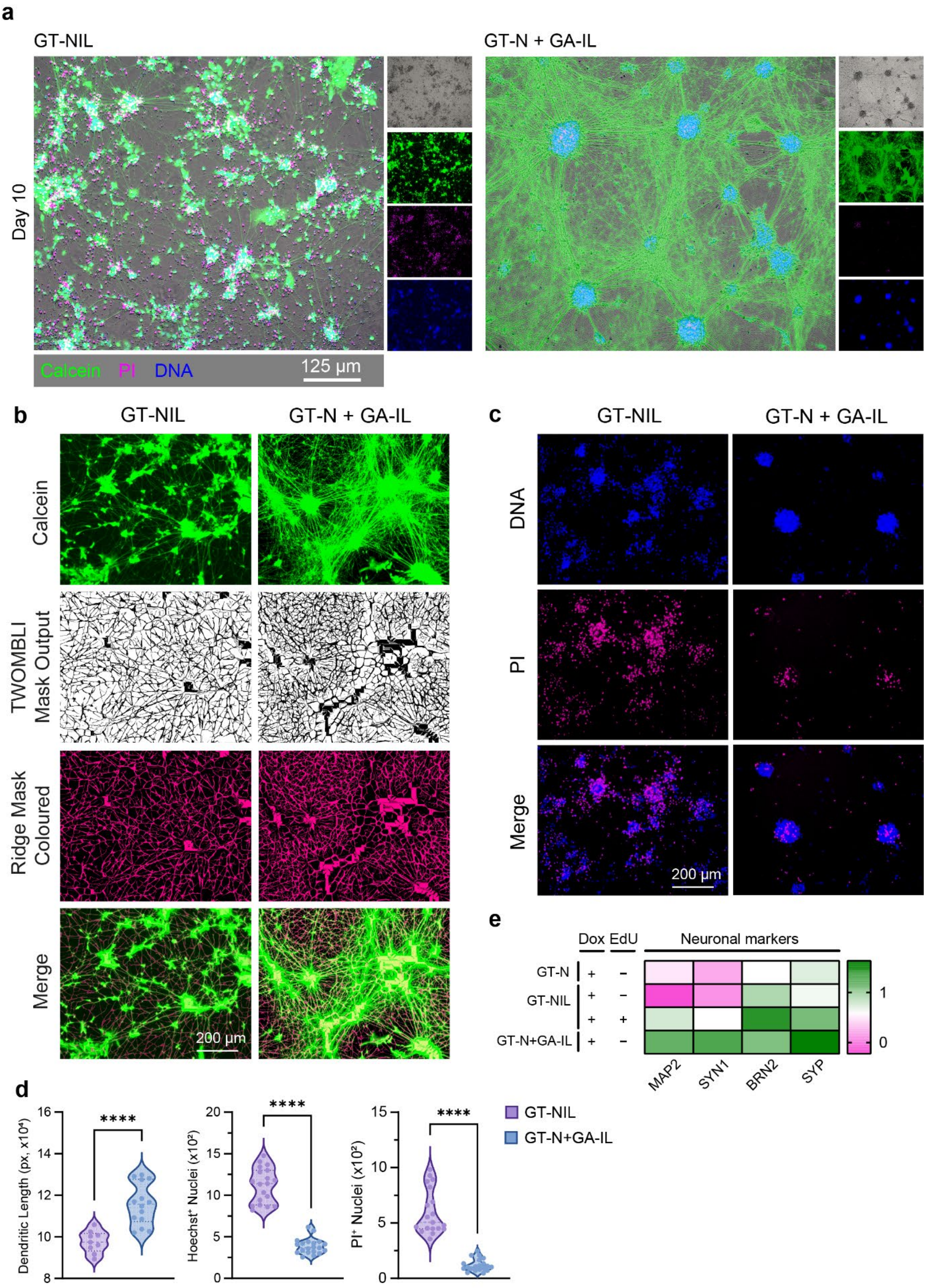

#### Supplementary Figure 9. Characterization of iMNs generated from GT-NIL and GT-N + GA-IL hiPSC lines

**(a)** Fluorescence/phase-contrast images on day 10 of doxycycline induction showing staining for live cells (Calcein AM, green), dead cells (propidium iodide, magenta) and DNA (Hoechst, blue).

**(b)** Representative fluorescence images of Calcein AM-stained live cells and quantified mask output of the mapped ridgelines.

**(c)** Representative fluorescence images of Hoechst-stained nuclei to visualize DNA, and propidium iodide $(PI)^+$ non-viable cells.

**(d)** Violin plots showing quantification of dendritic length (*left*), counts of $Hoechst^+$ nuclei (*middle*) and $PI^+$ nuclei (*right*) from images taken of cells as shown in panels **b** and **c**.

**(e)** Gene expression analysis of neuronal markers in the GT-N, GT-NIL, and GT-N + GA-IL lines after 10 days of doxycycline treatment. Values are normalized to RPL37A and shown relative to uninduced conditions ($log_{10}$-transformed). N=3 independent differentiations.

#### Supplementary Figure 10

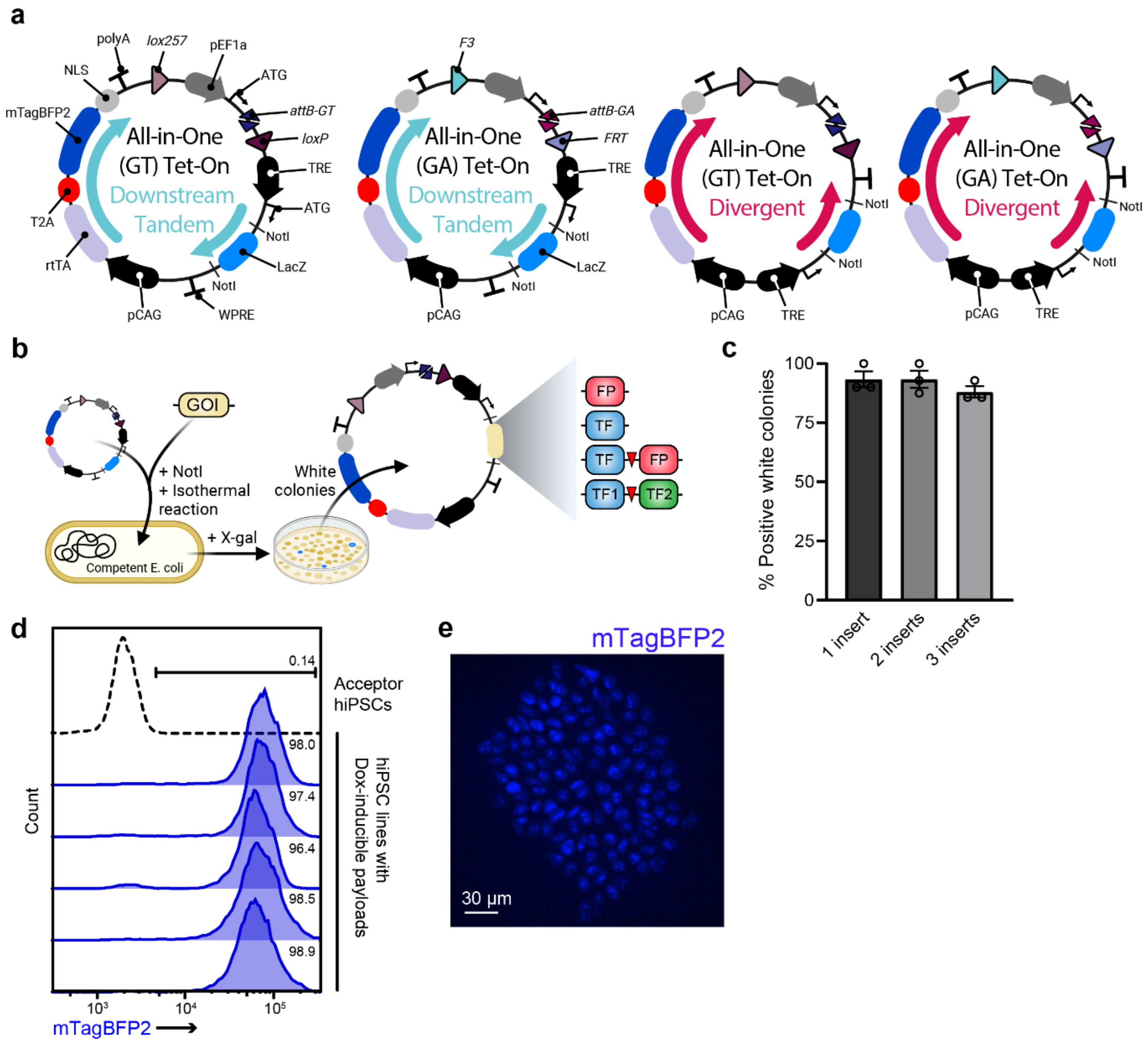

#### Supplementary Figure 10. Modular cloning platform for assembling downstream tandem and divergent Tet-ON 3G donor plasmids

**(a)** Schematic of GT and GA donor plasmids encoding all-in-one Tet-On 3G expression systems in either downstream tandem or divergent orientations.

**(b)** Overview of the cloning workflow, in which the *LacZ* cassette is replaced with one or more genes of interest (GOIs) via NotI digestion and isothermal assembly. IPTG/X-gal blue-white screening enables identification of clones containing the GOIs (white colonies).

**(c)** Mean percentages of white bacterial colonies that correctly assembled 1, 2, or 3 inserts into the downstream tandem donor plasmid, as confirmed by colony PCR. N=3 replicates; error bars, ±SEM.

**(d)** Flow cytometric analysis of mTagBFP2 expression in hiPSC lines with integrated downstream tandem Tet-On 3G GT donor plasmids containing various GOIs. Dashed line represents untransfected hiPSCs and values indicate the percentage of $mTagBFP2^{+}$ cells.

**(e)** Immunofluorescence image showing nuclear-localized mTagBFP2 expression in hiPSCs containing an integrated downstream tandem Tet-On 3G GT donor plasmid.

Supplementary Figure 11

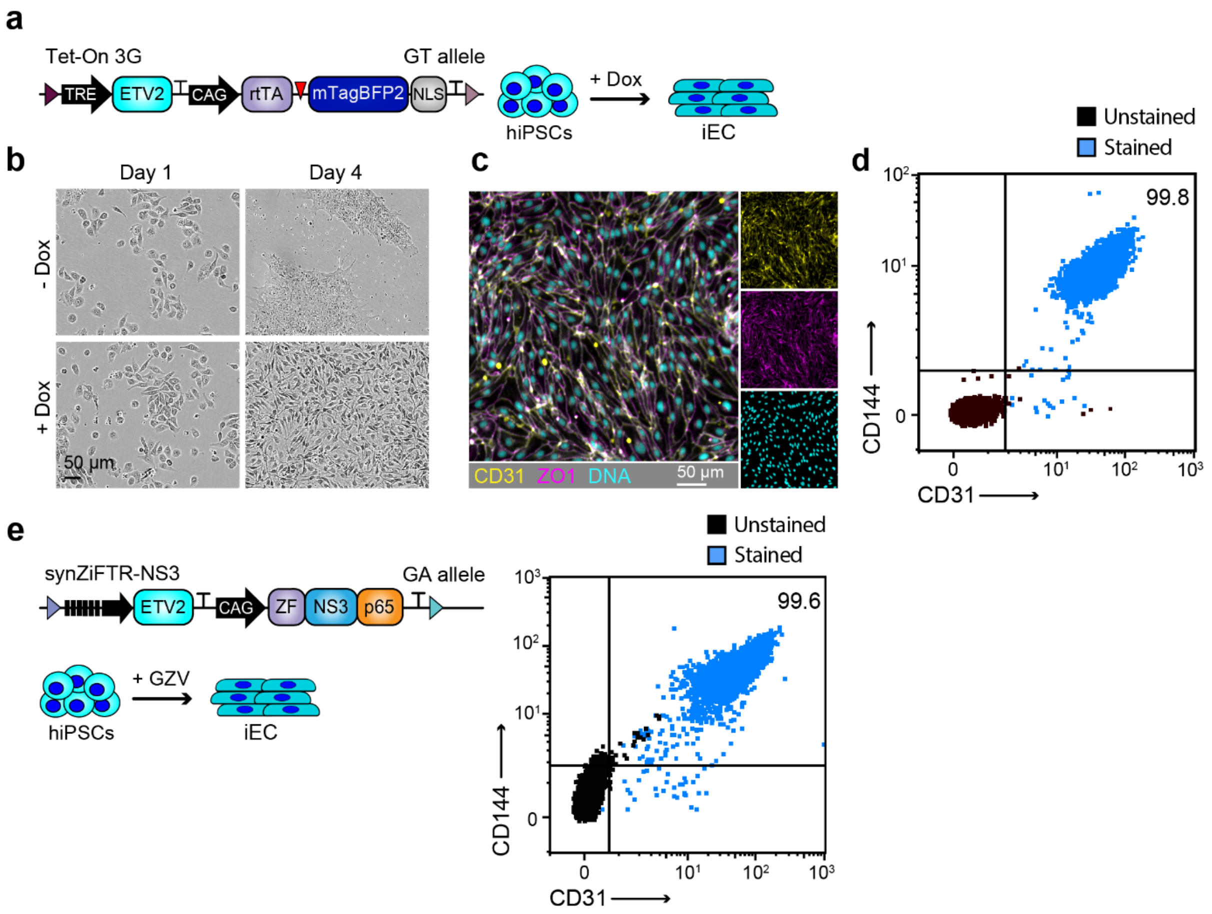


**Supplementary Figure 11. Characterization of iECs following *ETV2* overexpression using the Tet-On 3G system**

(**a**) Schematic of the downstream tandem configuration for a doxycycline-inducible *ETV2* expression cassette.

(**b**) Phase contrast images showing cell morphology at the indicated time points in the absence (-) or presence (+) of doxycycline.

(**c**) Immunofluorescence images of cells cultured with doxycycline for 4 days and stained for *CD31* (yellow), *ZO1* (magenta), and DNA (cyan).

(**d, e**) Flow cytometric analysis of endothelial markers *CD144* (VE-cadherin) and *CD31* (PECAM-1) on day 4 of *ETV2* induction using either the all-in-one Tet-On 3G (**d**) or synZiFTR-NS3 (**e**) inducible system. The iECs in (**e**) are derived from the dual-fate hiPSC line shown in **Figure 6**e, f.

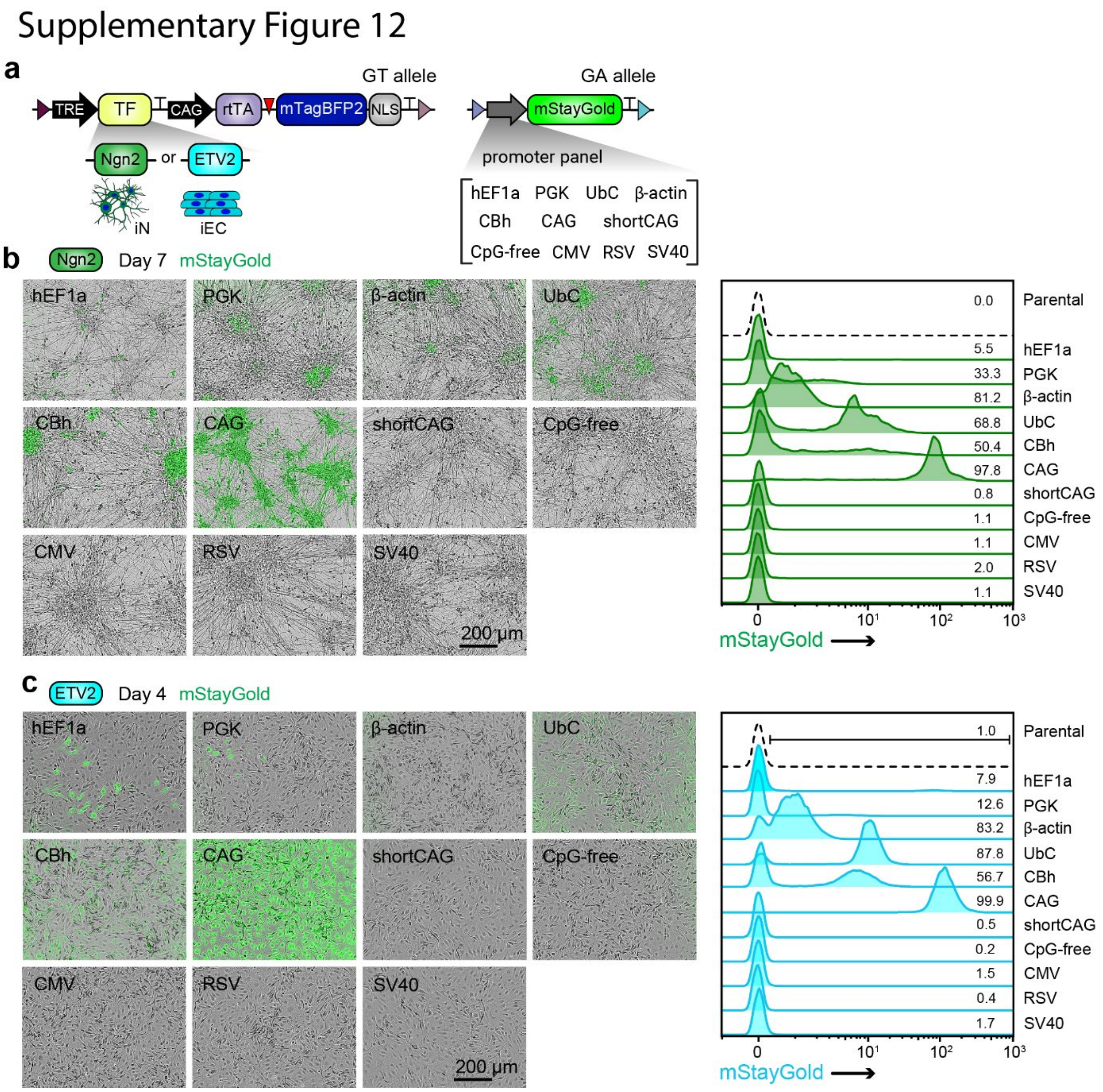


**Supplementary Figure 12. Evaluation of promoter activity in iNs and iECs using STRAIGHT-IN Dual**

(**a**) Schematic of the downstream tandem configuration for a doxycycline-inducible *Ngn2* or *ETV2* cassette integrated in the GT allele (*left*), and a schematic of the promoter panel driving mStayGold reporter expression integrated in the GA allele (*right*) of the *CLYBL* locus.

(**b, c**) Overlaid fluorescence and phase contrast images (*left*) and flow cytometric analysis of mStayGold expression (*right*) in cells cultured with doxycycline for 7 (**b**; *Ngn2*) or 4 days (**c**; *ETV2*). The promoter sequences assessed are indicated.

### Supplementary Figure 13

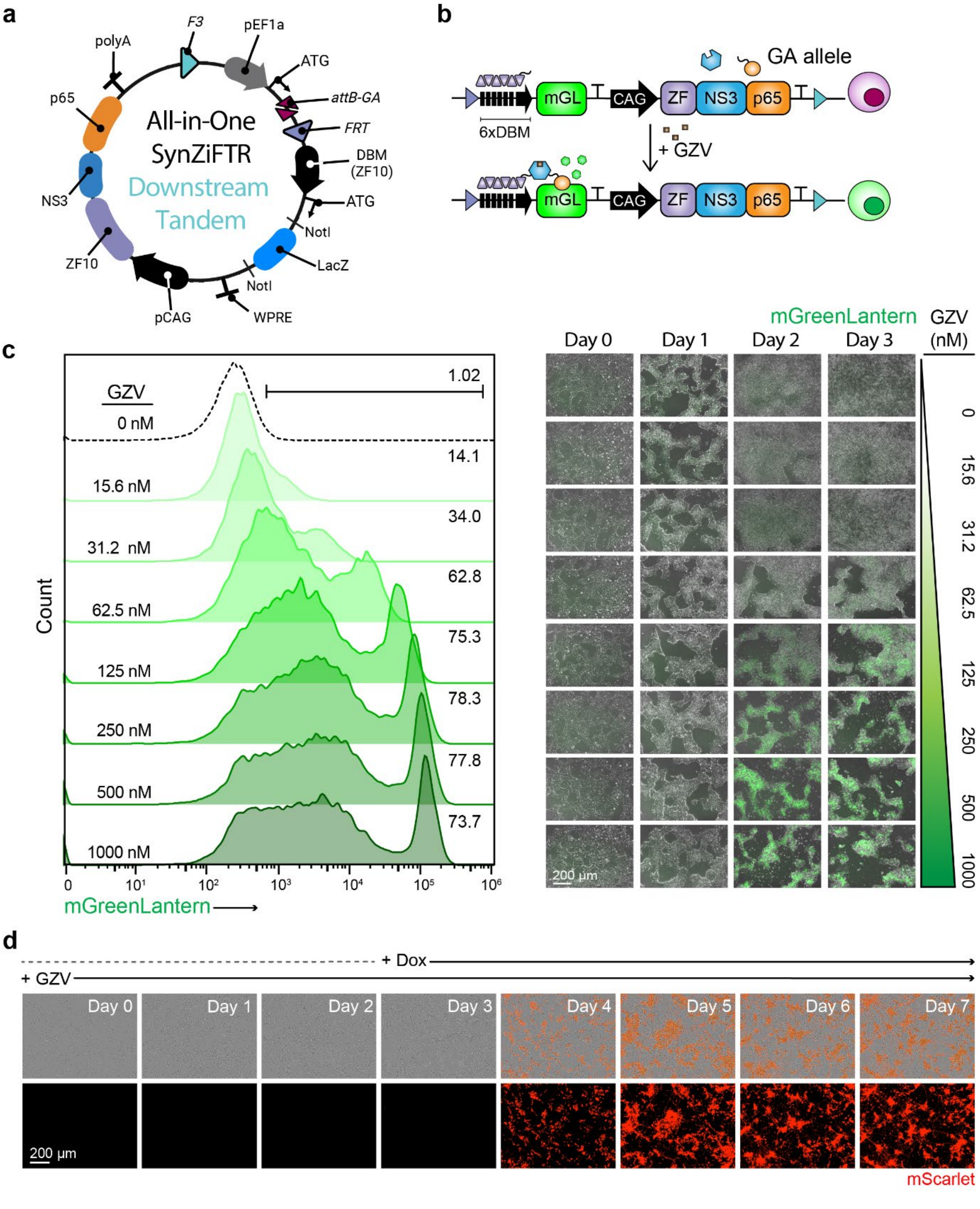

#### Supplementary Figure 13. Transgene overexpression in hiPSCs using the GZV-inducible synZiFTR system

**(a)** Schematic of the downstream tandem all-in-one GZV-inducible synZiFTR GA donor plasmid, in which the *LacZ* cassette is replaced with the gene of interest (GOI) via NotI digestion and isothermal assembly.

**(b)** Schematic of a downstream tandem synZiFTR-NS3 cassette driving GZV-inducible mGreenLantern expression, integrated in the GA allele.

**(c)** Flow cytometric analysis (*left*) and overlaid fluorescence and phase-contrast images (*right*) of hiPSCs containing the construct shown in (**b**) and cultured in the absence (dashed line) or presence (green) of increasing grazoprevir (GZV) concentrations over a 3-day period. Values in the plot indicate the percentage of cells expressing mGreenLantern.

**(d)** Timelapse fluorescence/phase-contrast images showing mScarlet expression in STRAIGHT-IN Dual hiPSCs containing the constructs shown in Figure 6C. Cells were cultured for 7 days with GZV added from day 0 and doxycycline from day 3.

Supplementary Figure 14

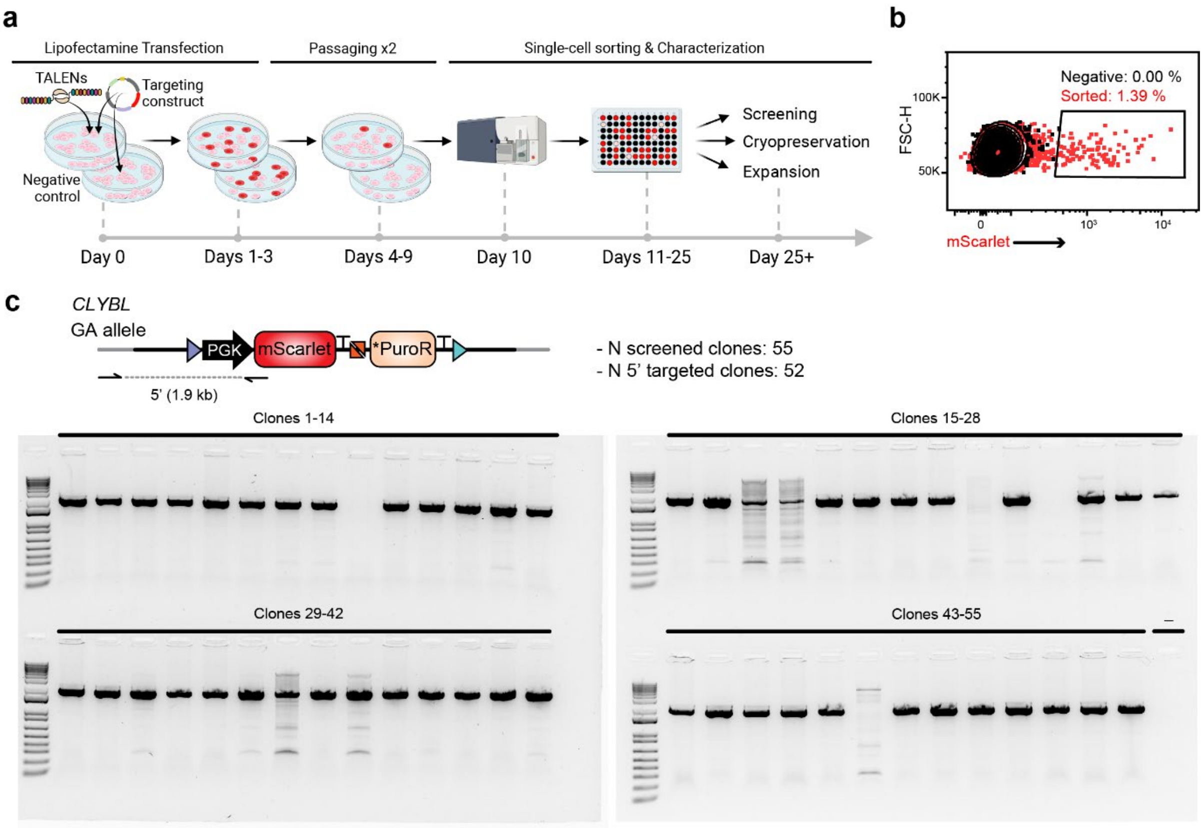


**Supplementary Figure 14. Targeting of Bxb1-GA landing pad into the *CLYBL* locus**

**(a)** Schematic of gene targeting workflow, lipofectamine-mediated co-transfection of the *CLYBL*-targeting construct and TALEN expression plasmids, followed by single-cell sorting and downstream clonal characterization.

**(b)** Flow cytometric analysis of hiPSCs expressing the mScarlet reporter from the GA landing pad.

**(c)** PCR screening results from 55 hiPSC clones using the primer pair indicated in the schematic to detect correct 5′ junction integration.

#### Supplementary Figure 15

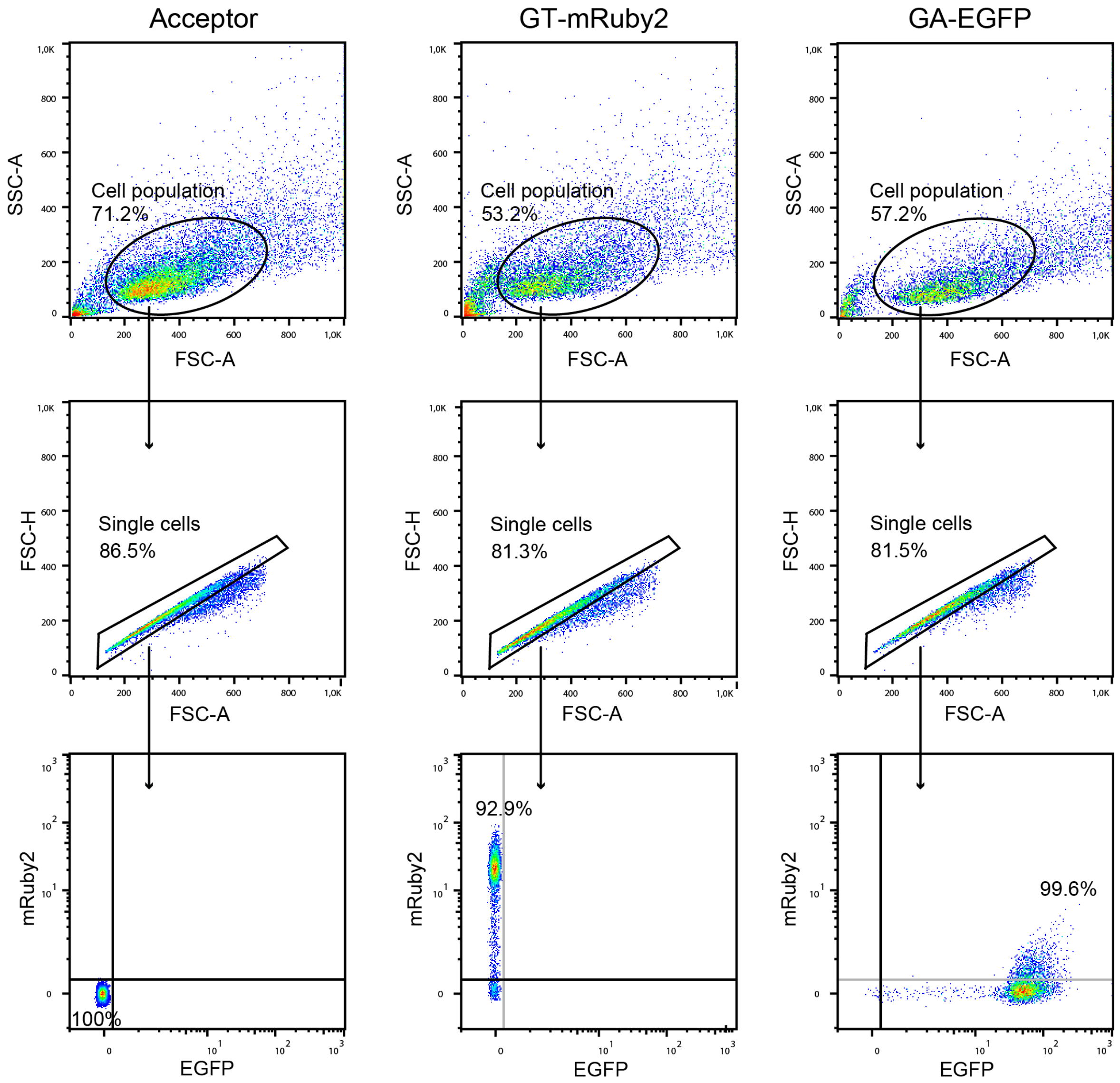


**Supplementary Figure 15. Examples of gating strategy used for flow cytometry analysis of hiPSCs**

Dot plots showing the gating workflow. Forward and side scatter was used to identify the cell population, with forward scatter area versus height used to distinguish single cells. Side scatter versus the appropriate fluorescence channel was used to gate *mRuby2*- or *EGFP*-positive hiPSCs.

#### Supplementary Table 1. Genotyping PCR Oligonucleotides

| Purpose | Sequence Forward Primer (5' - 3') | Sequence Reverse Primer (5' - 3') |
|---|---|---|
| *CLYBL_bxb1-GT* – 5' junction | AGATCATCCAGCCCTAGTCAAG | CGGTGGTGCAGATGAACTTC |
| *CLYBL_bxb1-GT* – 3' junction | AGCAAAGACCCCAACGAGAA | TGGAGCAGTGGATGACAACTT |
| *CLYBL_bxb1-GA* – 5' junction | AGATCATCCAGCCCTAGTCAAG | CCGTCCTCGAAGTTCATCAC |
| *CLYBL_bxb1-GA* – 3' junction | GACATCACCTCCCACAACGA | TGGAGCAGTGGATGACAACTT |

#### Supplementary Table 2. ddPCR Primer-Probe Sets

| Target Gene | Assay type | Primer/Probe | Sequence (5'- 3') | Fluorophore-Quencher | Source |
|---|---|---|---|---|---|
| *RPP30* | Copy number | Forward Primer<br>Reverse Primer<br>Probe | GATTTGGACCTGCGAGCG<br>GCGGCTGTCTCCACAAGT<br>CTGACCTGAAGGCTCT | HEX-ZEN-IBFQ | (Blanch-Asensio et al., 2024) |
| *EBFP2* | Copy number | Forward Primer<br>Reverse Primer<br>Probe | GCCGACAAGCAGAAGAACG<br>GGGTGTTCTGCTGGTAGTGG<br>AGATCCGCCACAACATCGAGG | FAM-ZEN-IBFQ | (Blanch-Asensio et al., 2024) |
| *mScarlet* | Copy number | Forward Primer<br>Reverse Primer<br>Probe | TGGCAACCTGACTTGTATCG<br>GTCCATCACTGTCCTTCACTATC<br>CGACAGGTGCTTCTCGATCTGCAT | FAM-ZEN-IBFQ | This study |
| *BleoR* | Copy number | Forward Primer<br>Reverse Primer<br>Probe | AGTTGACCAGTGCCGTTCC<br>CGAAGTCGTCCTCCACGAAG<br>AGCCGGTCGGTCCAGAAC | FAM-ZEN-IBFQ | (Blanch-Asensio et al., 2024) |
| *PuroR* | Copy number | Forward Primer<br>Reverse Primer<br>Probe | CGCCTTCCTGGAGACCTC<br>TTGCGGGTCATGCACCAG<br>CTCGGCTTCACCGTCACCG | FAM-ZEN-IBFQ | This study |
| *TurboGFP* | Copy number | Forward Primer<br>Reverse Primer<br>Probe | TGATGGGCTACGGCTTCTAC<br>CACCCGC+AT+CGAGAAGTACG<br>ATCACCTTGAAGTCGCCGATC | FAM-ZEN-IBFQ | This study |
| *AmpR* | Copy number | Forward Primer<br>Reverse Primer<br>Probe | TTTCCGTGTCGCCCTTATTCC<br>ATGTAACCCACTCGTGCACCC<br>TGCCTTCCTGTTTTTGCTCACCCA | FAM-ZEN-IBFQ | (Blanch-Asensio et al., 2024) |
| *pUC Ori* | Copy number | Forward Primer<br>Reverse Primer<br>Probe | CGATAAGTCGTGTCTTACCG<br>GCTTTCTCATAGCTCACGC<br>TGCACACAGCCCAGCTTG | FAM-ZEN-IBFQ | (Blanch-Asensio et al., 2024) |
| *mTagBFP2* | Copy number | Forward Primer<br>Reverse Primer<br>Probe | GGAGAACATGCACATGAAGCTGT<br>CCTGGGTAGCGGTCAGCA<br>AGGGCACCGTGGACAACC | FAM-ZEN-IBFQ | This study |
| *attP/attR* GT allele | Integration | Forward Primer<br>Reverse Primer<br>Reverse Primer<br>Probe<br>Probe | GCATTCTAGTTGTGGTTTGTCC<br>GCACTGGTCAACTTGGCATAT<br>CCTGCTTGCCGAATATCATG<br>CGTGGTTTGTCTGGTCAACCA<br>TCTCCGTCGTCAGGATCATCC | HEX-ZEN-IBFQ<br>HEX-ZEN-IBFQ | (Blanch-Asensio et al., 2023) |
| *attP/attR* GA allele | Integration | Forward Primer<br>Reverse Primer<br>Reverse Primer<br>Probe<br>Probe | CGTGGTTTGTCTGGTCAACCA<br>CGTGGGCTTGTACTCGGTAA<br>CTTAATTAACTAGTCGATGCCTGC<br>CGTGGTTTGTCTGGTCAACCA<br>ACTCCGTCGTCAGGATCATCC | HEX-ZEN-IBFQ<br>HEX-ZEN-IBFQ | This study |

*+ symbol indicates the following nucleotide is a locked nucleic acid*

#### Supplementary Table 3. Oligonucleotides used for NGS library

| Gene | Sequence (5' - 3') | Annealing temperature |
|---|---|---|
| EF1a_Fwd | GTTCAGAGTTCTACAGTCCGACGATCNNNNNNAGGAAAAGGGCCTTTCCGTCC | 67.2°C |
| Bactin_Fwd | GTTCAGAGTTCTACAGTCCGACGATCNNNNNNCCTCCGACCAGTGTTTGCCT | 71.1°C |
| CpGfree_Fwd | GTTCAGAGTTCTACAGTCCGACGATCNNNNNNAGTACTCCCTCTCAAAAGCTGGC | 71.1°C |
| PGK_Fwd | GTTCAGAGTTCTACAGTCCGACGATCNNNNNNCTCCGCCCCTAAGTCGGGAA | 67.2°C |
| RSV_Fwd | GTTCAGAGTTCTACAGTCCGACGATCNNNNNNGGTAACGATGAGTTAGCAACATGCC | 67.2°C |
| UbC_Fwd | GTTCAGAGTTCTACAGTCCGACGATCNNNNNNGTGAGGCGTCAGTTTCTTTGGTCG | 67.2°C |
| SV40_Fwd | GTTCAGAGTTCTACAGTCCGACGATCNNNNNNGTTAATTAAGTACTTACTGCAGGCAGAA | 67.2°C |
| CMV_Fwd | GTTCAGAGTTCTACAGTCCGACGATCNNNNNNGCACCAAAATCAACGGGACTTTCC | 67.2°C |
| CAG_Fwd | GTTCAGAGTTCTACAGTCCGACGATCNNNNNNGTTCGGCTTCTGGCGTGTGA | 67.2°C |
| CBh_Fwd | GTTCAGAGTTCTACAGTCCGACGATCNNNNNNAAGAGGTAAGGGTTTAAGGGATGGT | 67.2°C |
| Generic_Rev | TTCCTTGGCACCCGAGAATTCCACGTGGAACCAGTTCTTCAGGC | 67.2 or 71.1°C |

#### Supplementary Table 4. qPCR Oligonucleotide Pairs

| Gene | Sequence Forward Primer (5' - 3') | Sequence Reverse Primer (5' - 3') |
|---|---|---|
| *RPL37A* | GTGGTTCCTGCATGAAGACAGTG | TTCTGATGGCGGACTTTACCG |
| *OCT4* | GTGGAGGAAGCTGACAACAA | ATTCTCCAGGTTGCCTCTCA |
| *NANOG* | AGCAGATGCAAGAACTCTCCAA | TGAGGCCTTCTGCGTCACAC |
| *KLF4* | ATAGCCTAAATGATGGTGCTTGG | AACTTTGGCTTCCTTGTTTGG |
| *SOX2* | GCTACAGCATGATGCAGGACCA | TCTGCGAGCTGGTCATGGAGTT |
| *TUBB3* | CAACCAGATCGGGGCCAAGTT | CCGAGTCGCCCACGTAGTT |
| *MAP2* | AGACTGCAGCTCTGCCTTTAG | AGGCTGTAAGTAAATCTTCCTCC |
| *SYN1* | CCCTGGGTGTTTGCCCAGAT | ACCACGGGGTACGTTGTACT |
| *BRN2* | ACCCGCTTTATCGAAGGCAA | CCTCCATAACCTCCCCCAGA |
| *GRIA4* | GGCCAGGGAATTGACATGGA | AACCAACCTTTCTAGGTCCTGTG |
| *VGLUT2* | GTAGACTGGCAACCACCTCC | CCATTCCAAAGCTTCCGTAGAC |
| *SYP* | ACCTCGGGACTCAACACCTCGG | GAACCACAGGTTGCCGACCCAG |
| *Ngn2* | TATGCACCTCACCTCCCCATAG | GAAGGGAGGAGGGCTCGACT |
| *ISL1* | AAGGTGGAGCTGCATTGGTTTG | TAAACCAGCTACAGGACAGGCC |
| *CHAT* | CTCAGCTACAAGGCCCTGCT | ACCAGCGTGTCCTGGGTATG |
| *FOXA2* | GGAGCGGTGAAGATGGAA | TACGTGTTCATGCCGTTCAT |
| *PDX1* | GATGAAGTCTACCAAAGCTCACG | GTTCAACATGACAGCCAGCTC |
| *SOX17* | CGCACGGAATTTGAACAGTA | GGATCAGGGACCTGTCACAC |
| *ACTA2* | GTGATCACCATCGGAAATGAA | TCATGATGCTGTTGTAGGTGGT |
| *VIM* | AGTCCACTGAGTACCGGAGAC | CATTTCACGCATCTGGCGTTC |
| *T* | TGTTTATCCATGCTGCAATCC | CCGTTGCTCACAGACCACAG |
| *PAX6* | CGAGATTTCAGAGCCCCATA | AAGACACCACCGAGCTGATT |
| *LHX3* | ACAGACACTGGCACAGCAAG | AGCAGTGCAGATGGTACACG |
| *HB9* | GCCTAAGATGCCCGACTTCAAC | CGCGACAGGTACTTGTTGAGCT |
| *VAChAT* | GCTGTTTGCTTCCAAGGCTATCC | GAAGGCGAACAGGACTGTAGAG |
| *LHX4* | TGGCGGACAGGTGCTTCTCCA | AGGTGGTAGACAAAGTCCTGGG |
| *CDH5* | CCTACCAGCCCAAAGTGTGT | TGTCCTTGTCTATTGCGGAGA |
| *CD31* | ATGCCGTGGAAAGCAGATAC | CTGTTCTTCTCGGAACATGGA |
| *TIE2* | ACGACCATGACGGCGAATG | CGGCAGCCTGATATGCCTG |
| *FLK1* | CGGACAGTGGTATGGTTCTTG | GCCACAGACTCCCTGCTTT |
| *TAL1* | CCAACAATCGAGTGAAGAGGA | CCGGCTGTTGGTGAAGATAC |
